# Improving the efficiency of single-driver ecological experiments with Bayesian optimal experimental design

**DOI:** 10.64898/2026.05.28.728579

**Authors:** Ravi Ranjan, Mridul K. Thomas

## Abstract

1. Ecological experiments often characterize species responses to environmental drivers by estimating parameters of well-known nonlinear functions. However, the standard experimental designs used for these experiments waste precious experimental resources by making measurements at uninformative driver levels.
2. Classical methods to optimize experimental designs require the parameter values we intend to estimate – circularity that undermines the usefulness of optimization. Bayesian Optimal Experimental Design (BOED) solves this problem by using prior distributions of the parameters to calculate designs that optimize properties of the posterior distribution. Thus, they circumvent the parameter dependence and result in robust, efficient experimental designs.
3. Here, we develop and evaluate Bayesian optimal designs for four commonly used nonlinear drivers measuring per-capita growth rate against: nutrients/food (Monod or Holling type 2 function), light (Eilers-Peeters function), temperature (Norberg function) and toxins (log-logistic function).
4. We show using simulations that Bayesian optimal designs consistently outperform standard uniform designs in terms of parameter estimation and prediction accuracy, especially at low sample sizes. For some functions, Bayesian designs with 5 data points outperformed uniform designs with 15 data points. BOED can therefore allow us to allocate scarce experimental resources more efficiently. We provide detailed explanations and code to enable readers to apply these methods. We also provide rules of thumb that would improve experimental efficiency even without following the entire BOED procedure.

## 1. Introduction

When measuring the effect of a continuous driver, gradient designs paired with a regression analysis are the standard tool (Collins et al., 2022; Cottingham et al., 2005; Schweiger et al., 2026). Unlike categorical designs that ask *if* a response changes between two levels (or more in rare cases) of the driver, a gradient design goes further and asks: *how* does the response change as a continuous function of the driver? This additional information can often be incorporated into theoretical models or used to make accurate predictions. Since most environmental drivers like nutrients, light, temperature and toxin concentrations are continuous, single-driver gradient experiments are especially common in ecology (Cottingham et al., 2005).

Despite their commonness, gradient experiments are almost always performed inefficiently. Gradient experiments typically have an approximately uniform design in ecology, where replicated measurements are made at equally spaced levels across the range of the driver (Cottingham et al., 2005). However, all levels across a gradient are not equally informative. The model fit, especially for non-linear models, is more sensitive to some levels than others. We can see this in simulations of a Monod growth experiment, where per-capita growth rate is measured at different nutrient concentrations (Fig. 1a). In Fig. 1b, we show the *generalized leverage* of the Monod curve in Fig. 1a. Generalized leverage is a measure of relative influence of a measurement on the regression parameter estimates (Burch et al., 2012). Measurements in regions with high leverage values have a large influence on parameter estimates. We considered two designs for our Monod simulation: a uniform design where 5 points are spaced uniformly across the nutrient range and a custom design where we placed the 5 points only at the two local maxima of the generalized leverage curve. We simulated 100,000 datasets using each of these designs with a specific maximum growth rate *μ*_max_ and half-saturation constant *K*, and estimated the parameters from both simulated datasets. The custom designs result in a distribution of estimates with both lower variance and bias than the uniform design (Fig. 1c, d). Thus, uniform designs are often inefficient because they waste experimental effort on low-leverage levels.

**Figure 1:**
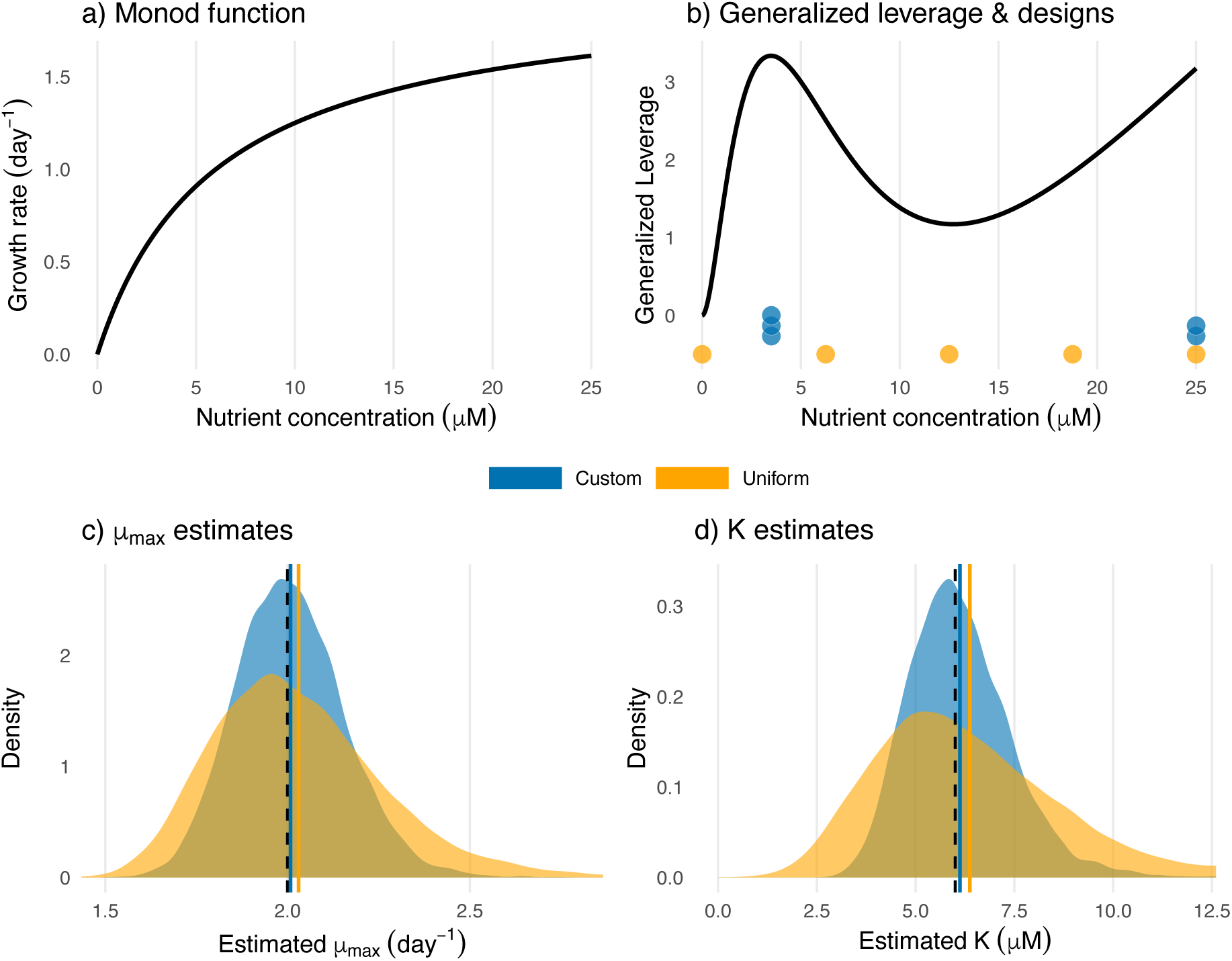
All levels in a regression are not equally informative. a) The Monod function describes per-capita growth rate against nutrient concentration. b) Generalized leverage measures the relative inffuence of a measurement at a point on the parameter estimates. Two designs are considered: a uniform design (orange) and a custom design (blue) where experimental units are placed at the two local maxima of the leverage curve. c) and d) show 100,000 maximum growth rate μ_max_ and half-saturation constant K estimates from simulated datasets based on the uniform and custom designs. For both parameters, the mean of the custom design estimates (blue solid line) is closer to the true value (black dashed line; μ_max_ = 2 day^−1^, K = 6 μM) than the mean of uniform design estimates (orange solid line).

Optimal experimental design theory uses the Fisher information matrix of the focal function to optimize the levels at which experimental units are placed (Berger & Wong, 2009; Huan et al., 2024; Thomas & Ranjan, 2024). It strategically places experimental units to achieve a goal such as minimizing parameter estimation error or prediction error. The resulting designs are optimal with respect to the specified goal and are referred to as classical optimal designs. Several barriers have prevented the widespread adoption of optimal design methods. First, classical optimal designs change based on the function being fitted and therefore require prior knowledge of the function. Thus, classical optimal designs cannot be used for exploratory experiments where the response is completely unknown. However, this is not an obstacle to many ecological and eco-physiological experiments – such as how population growth rate responds to nutrients, light, temperature and toxins – that have well-established functions (Eilers C Peeters, 1988; Monod, 1950; Norberg, 2004; Ritz, 2010). Second, in non-linear regressions, classical optimal designs depend on the specific parameter values of the function being fitted. This creates a Catch-22 where the parameters cannot be obtained without the experiment, and the experiment cannot be designed without the parameters. However, we can break this Catch-22 by using prior distributions of possible values to find a Bayesian optimal design (Chaloner & Verdinelli, 1995). For many single-driver ecological experiments, past experiments provide plenty of data to develop reasonable Bayesian priors. As clear solutions exist to both these barriers, single-driver regression experiments present an avenue where optimal designs can rapidly reduce experimental effort and resources used. Since these experiments are common across taxa and are often a part of bigger projects, the benefits of making them more efficient and faster through optimal designs could be massive. Especially with the rise of robot-and AI-assisted experiments that can take place continuously, efficiencies compound through time, leading to more rapid discovery.

Given a prior distribution of the parameters, Bayesian Optimal Experimental Design (BOED) is a framework to find a design that optimizes criteria related to the posterior distribution. A commonly used criterion focused on parameter inference is the expected Shannon Information Gain (SIG) between the prior and the posterior. Other criteria focused on prediction or model discrimination also exist (Huan et al., 2024; Ryan et al., 2016). Thus, BOED uses both the prior and the posterior to calculate optimal designs that are robust across possible parameter values and efficient even at small sample sizes.

**Here, we develop Bayesian optimal designs for four common single-driver experiments in ecology** measuring how population growth (or performance) changes with: 1) nutrient concentration (Monod function), 2) light (Eilers-Peeters function), 3) temperature (Norberg function) and 4) toxin concentration (log-logistic function, a type of dose-response curve). These functions apply broadly across the tree of life: the nutrient and toxin functions apply to all organisms, the light function applies to all photosynthetic organisms and the temperature function applies to all ectotherms. All are non-linear, but their shapes range from monotonically increasing but saturating for nutrients, to unimodal curves for temperature and light (with opposite skewness) and an S-shaped curve for toxins. While we select parameter values that reflect phytoplankton experiments, these can easily be changed to reflect other taxa. We compare optimal versus uniform designs and use simulations to evaluate their performance in estimating parameters and predictive ability. We present the workflow for calculating these designs along with associated code. We also present rules of thumb to make these experiments more efficient even for readers who do not wish to apply the BOED approach.

## 2. Optimal experimental design

### 2.1 Classical optimal designs

We consider a non-linear regression where the function *g* relates the value of focal environmental variable *x_i_* to the corresponding specific growth rate measurement *μ_i_* with observation error *ε_i_*.

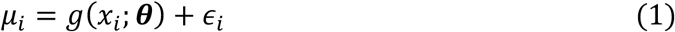

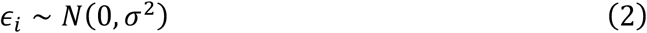

The function *g* specifies the mean growth rate at an experimental unit with environmental driver value *x_i_* (*i* = 1…*n*). The unknown parameter vector ***θ*** is of length *p*. The observation errors are normally distributed with mean 0 and standard deviation *σ*. While errors typically tend to be proportional to growth rates in such experiments, we do not explore that here. The overall parameter vector is thus ***ψ****^T^* = (***θ*^T^**, *σ*). The set of experimental units is the design ***δ*** = (*x*_1_…*x_n_*)^T^. The set of all operationally possible experimental units *x_i_* is the *design space*. The *design matrix* ***X*** (*n* × *p*) captures how growth rate at each experimental unit in the design changes with the parameter values. It can be defined as:

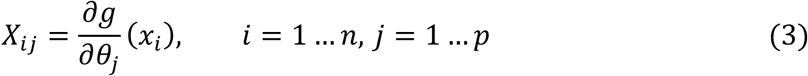

A design matrix element *X_i_*_j_ is the sensitivity (*∂g*/*∂θ*_j_) of a predicted growth rate *g*(*x_i_*) to changes in parameter *θ*_j_. The design matrix can be used to calculate the Fisher information matrix ***I***(***ψ***; ***δ***), which measures the amount of information that the growth rate data ***μ*** carries about the parameters ***θ***:

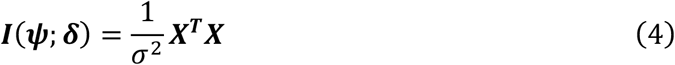

Classical optimal designs maximize different characteristics of the information matrix, each corresponding to a specific objective such as parameter estimation or prediction ability. However, in the case of non-linear regressions, the Fisher information matrix depends on the parameters of the model. This creates the aforementioned Catch-22: an optimal design cannot be calculated without specific parameter values and the parameter values cannot be found without conducting the experiment.

### 2.2 Bayesian Optimal Experimental Design (BOED)

To break the Catch-22, Bayesian Optimal Experimental Design (BOED) uses prior distributions of parameters. Given a prior distribution, BOED seeks to find an optimal design that results in the ‘best’ posterior (Chaloner & Verdinelli, 1995; Ryan et al., 2016), defined using different criteria. For example, a useful BOED criterion is the Shannon Information Gain (SIG, *u*_SIG_) between the prior (*π*(***θ***)) and the posterior (*π*(***θ***|***μ***, ***δ***)):

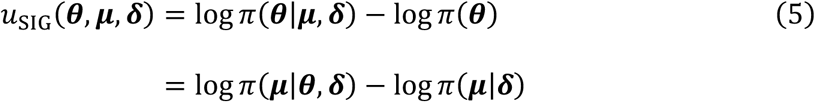

Intuitively, SIG is the difference between our knowledge before and after the experiment. If the prior (*π*(***θ***)) is flat and the experimental data results in a sharp posterior (*π*(***θ***|***μ***, ***δ***)), there is a large reduction in uncertainty and the SIG value is high.

Calculating SIG requires knowledge of the dataset (***μ***), which is unknown without doing the experiment. Therefore, instead of *u*_SIG_, the *expectation* of the SIG value (*U*_SIG_) is calculated over *all possible future* datasets for a given design. The probability of obtaining a specific dataset is the marginal likelihood (*π*(***μ***|***δ***)), calculated by integrating the likelihood (*π*(***μ***|***θ***, ***δ***)) over the prior distribution (*π*(***θ***)). Thus, the expected SIG utility *U*_SIG_(***δ***) of a design ***δ*** can be written as:

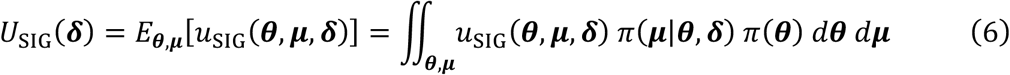

The integral is analytically intractable and is evaluated numerically using Monte Carlo techniques (see Section 1, Supplementary Information (SI) for details). Since *U*_SIG_ must be evaluated for each candidate design, finding the best design is computationally expensive especially when the dimensionality of the experiment (product of number of experimental units *n* and number of variables *v*) is high. Therefore, we use a two-phase stochastic optimization algorithm called Approximate Coordinate Exchange (ACE; overview in Section 2, SI) to calculate these designs (Overstall & Woods, 2017). ACE can calculate Bayesian optimal designs for non-linear regressions in a reasonable amount of time (Overstall & Woods, 2017). The ACE algorithm is implemented in the R package *acebayes* (Overstall et al., 2020).

### 2.3 Four focal models

In this study, we develop optimal designs for four common non-linear functions measuring growth (or performance) vs: 1) nutrient concentration (Monod function; Fig. 2a; Monod, 1950), 2) light (Eilers-Peeters function; Fig. 3a; Eilers & Peeters, 1988), 3) temperature (Norberg function; Fig. 4a; Baker et al., 2016; Norberg, 2004) and 4) toxin concentration (log-logistic function; Fig. 5a; Ritz, 2010).

**Figure 2:**
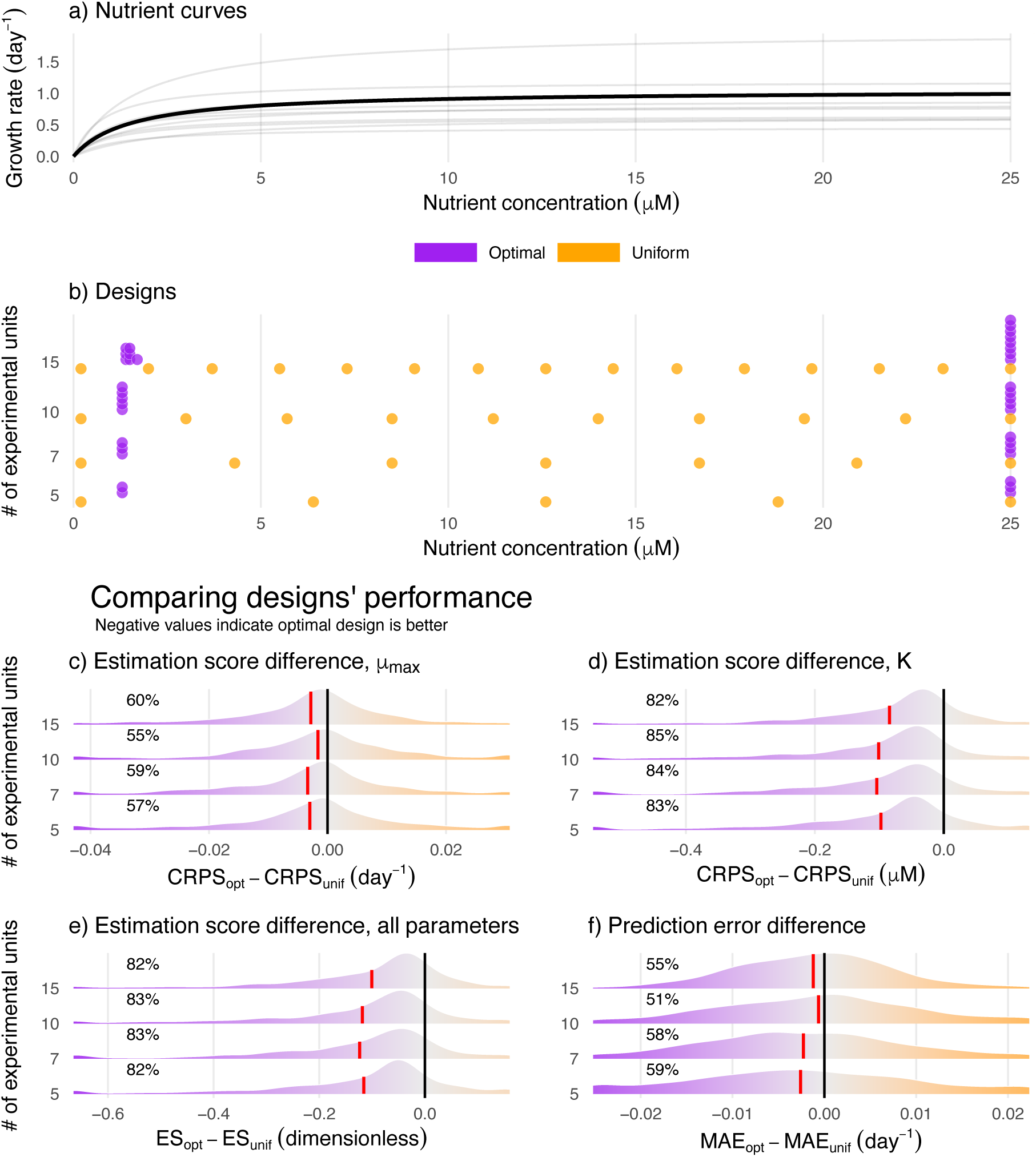
For the Monod function, optimal designs (purple) generally outperform uniform designs (orange). a) The central curve (thick black) along with 10 curves (grey) with parameters drawn from the prior. b) Optimal and uniform designs for sample sizes of 5, 7, 10 and 15 experimental units. c-f) Ridgeline plots showing the distribution of differences in the CRPS, Energy Score (ES) and aggregate prediction error of the optimal and uniform designs, across 1000 paired simulations at each sample size. The distributions have been trimmed such that values in the bottom and top 2 percentiles of all 4000 samples are pooled together at the edge of the plots. The black bar shows a difference of zero and the red bar denotes the mean difference between the scores. Negative values (to the left of the black bar) indicate simulations where the optimal design performed better and are shown in purple. The percentage of simulations where optimal design performs better is shown on the left of the plots. Optimal designs are consistently better for estimating K and joint estimation of all parameters. Optimal designs are slightly better for estimating μ_max_ and aggregate prediction error.

**Figure 3:**
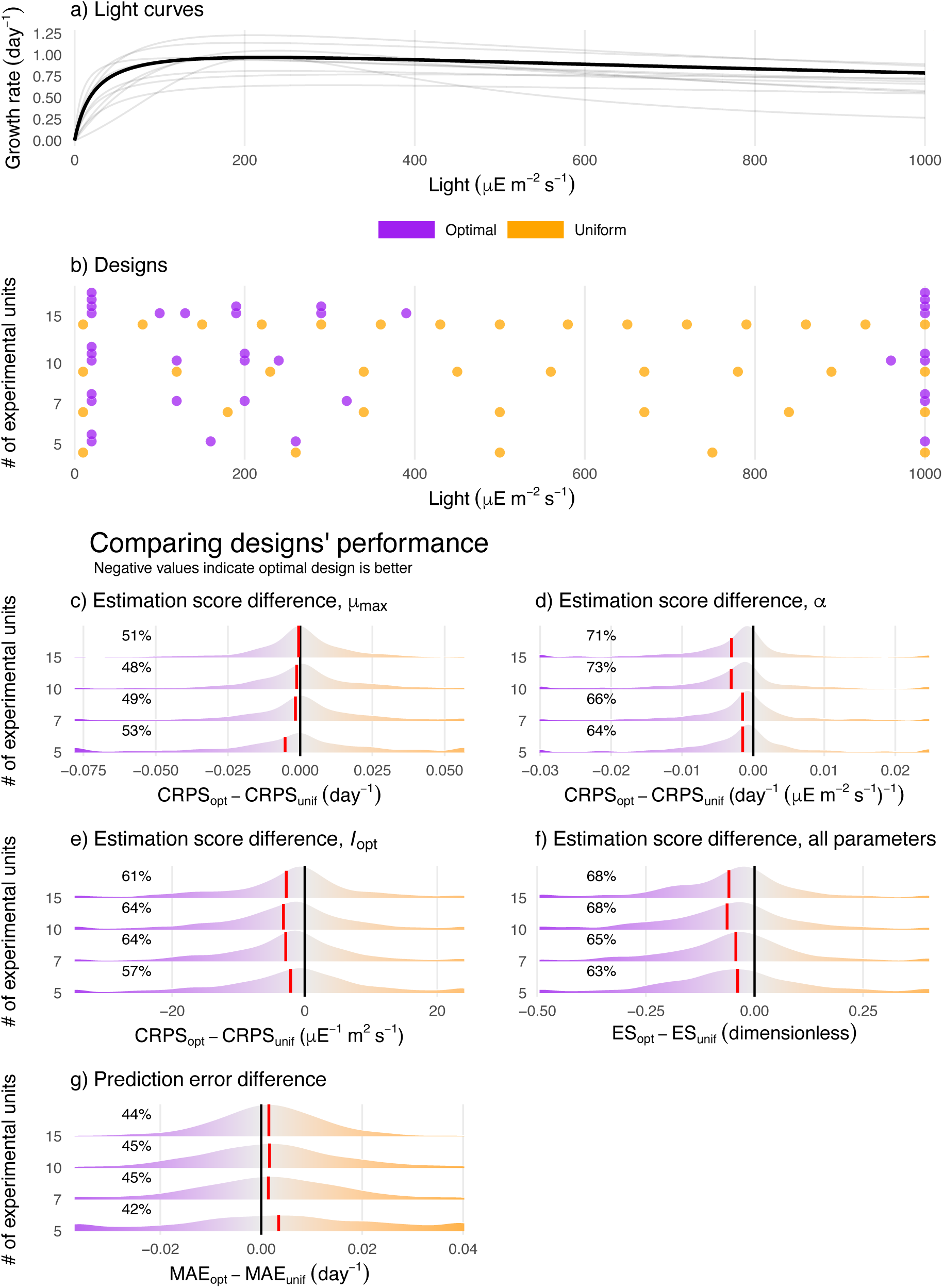
For the Eilers-Peeters function, optimal designs (purple) generally outperform uniform designs (orange). a) The central curve (thick black) along with 10 curves (grey) with parameters drawn from the prior. b) Optimal and uniform designs with 5, 7, 10 and 15 experimental units. c-g) Ridgeline plots showing the distribution of differences in the CRPS, Energy Score (ES) and aggregate prediction error of the optimal and uniform designs, across 1000 paired simulations at each sample size. The distributions have been trimmed such that values in the bottom and top 2 percentiles of all 4000 samples are pooled together at the edge of the plots. The black bar shows a difference of zero and the red bar denotes the mean difference between the scores. Negative values (to the left of the black bar) indicate simulations where the optimal design performed better and are shown in purple. The percentage of simulations where optimal design performs better is shown on the left of the plots. Optimal designs are substantially better for estimating α and joint estimation of all parameters. Optimal designs are slightly better for estimating I_opt_, μ_max_ and aggregate prediction errors.

**Figure 4:**
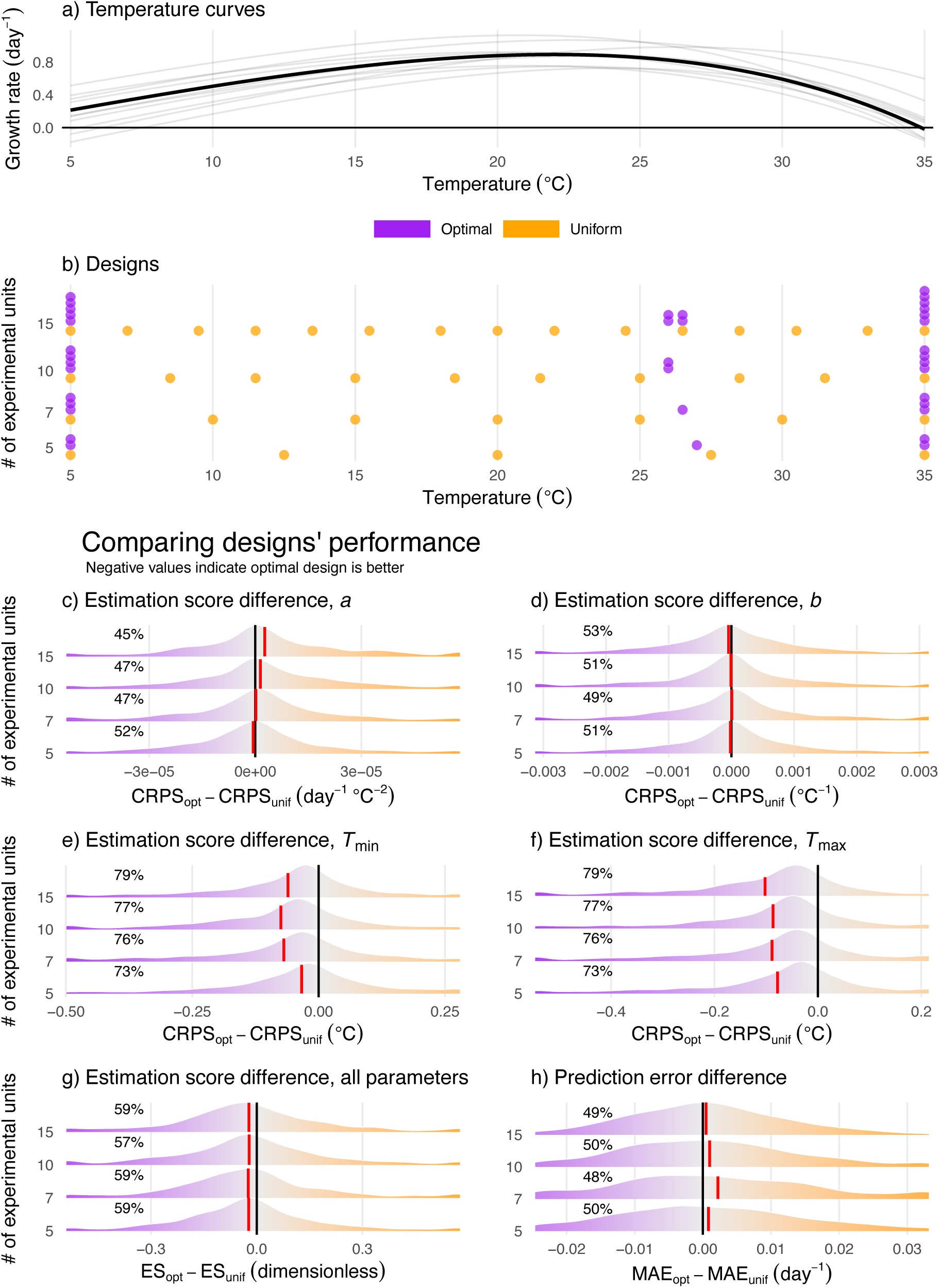
For the Norberg function, optimal designs (purple) generally outperform uniform designs (orange). For the temperature function alone, we enforced correlations between the parameters via the prior, to restrict ourselves to realistic curve shapes. a) The central curve (thick black) along with 10 curves (grey) with parameters drawn from the prior along with. b) Optimal and uniform designs with 5, 7, 10 and 15 experimental units. c-h) Ridgeline plots showing the distribution of differences in the CRPS, Energy Score (ES), and aggregate prediction error of the optimal and uniform designs, across 1000 paired simulations at each sample size. The distributions have been trimmed such that values in the bottom and top 2 percentiles of all 4000 samples are pooled together at the edge of the plots. The black bar shows a difference of zero and the red bar denotes the mean difference between the scores. Negative values (to the left of the black bar) indicate simulations where the optimal design performed better and are shown in purple. The percentage of simulations where optimal design performs better is shown on the left of the plots. Optimal designs are substantially better for estimating T_max_, T_min_ and joint estimation of all parameters. Optimal designs are similar to uniform designs for estimating b and slightly worse for estimating a.

**Figure 5:**
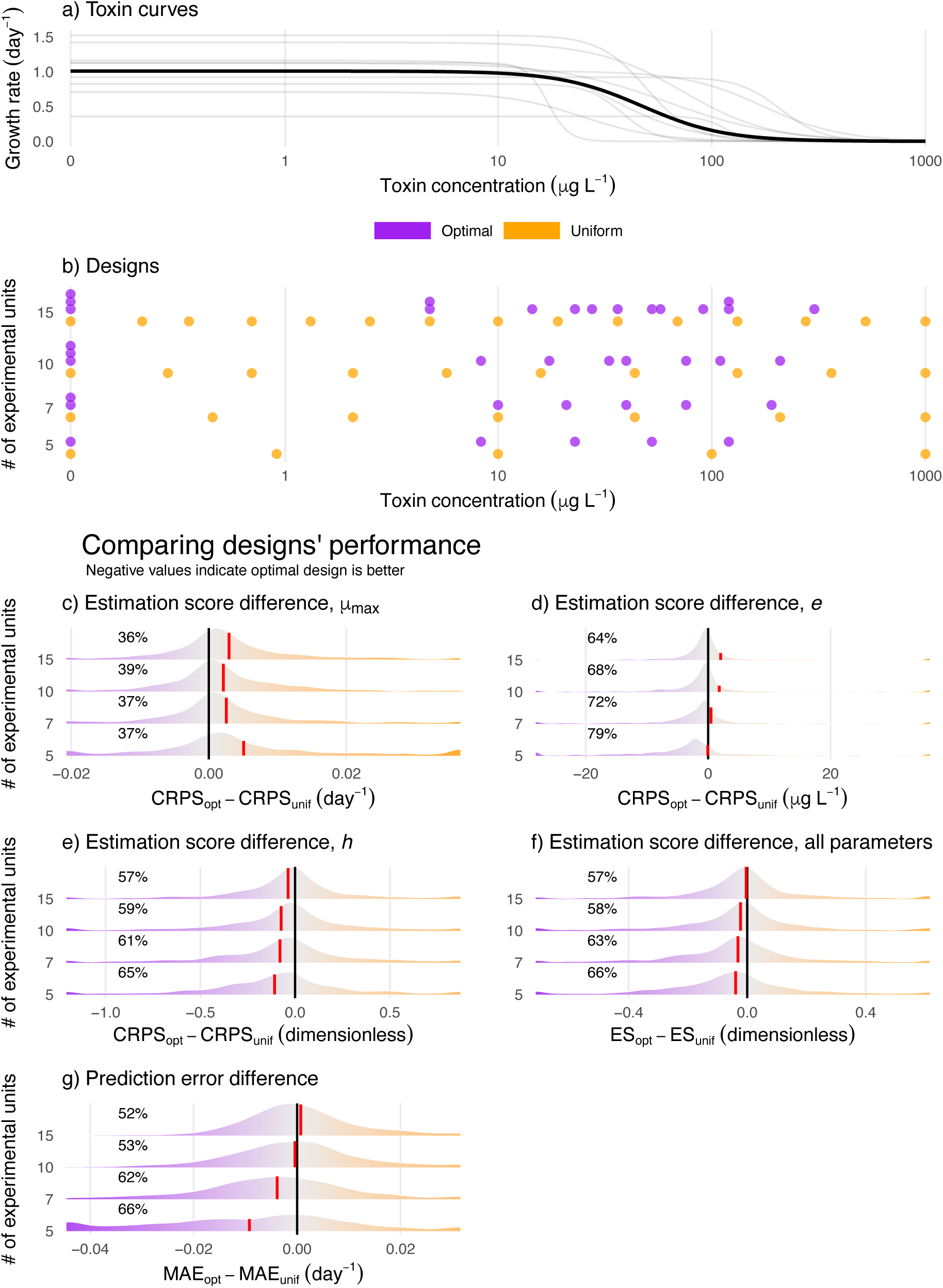
For the log-logistic function, optimal designs (purple) generally outperform uniform designs (orange). Note that panels a and b are plotted on a pseudo-logarithmic scale, which transitions from linear near zero to logarithmic around 0.1. a) The central curve (thick black) along with 10 curves (grey) with parameters drawn from the prior. b) Optimal and uniform designs with 5, 7, 10 and 15 experimental units. c-g) Ridgeline plots showing the distribution of differences in the CRPS, Energy Score (ES) and aggregate prediction error of the optimal and uniform designs, across 1000 paired simulations at each sample size. The distributions have been trimmed such that values in the bottom and top 2 percentiles of all 4000 samples are pooled together at the edge of the plots. The black bar shows a difference of zero and the red bar denotes the mean difference between the scores. Negative values (to the left of the black bar) indicate simulations where the optimal design performed better and are shown in purple. The percentage of simulations where optimal design performs better is shown on the left of the plots. Optimal designs are substantially better for estimating e, ℎ, joint estimation of all parameters and aggregate prediction error. The positive mean CRPS differences for e are a result of a few extreme EC50 true values resulting from the prior. Optimal designs are worse for estimating μ_max_.

The Monod function describes how per-capita growth rate (*μ*(*S*)) increases as a saturating function of nutrient concentration, *S* (Fig. 2a):

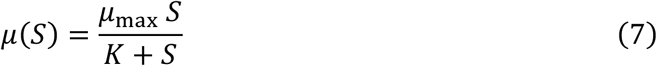

Here, *μ*_max_ is the maximum growth rate and *K* is the half-saturation constant. The Monod function is typically used to measure growth for microbes, although the function is the same as the Holling Type II functional response.

The Eilers-Peeters function describes per-capita growth rate (*μ*(*I*)) of phytoplankton populations against incident light intensity or irradiance, *I* (Fig. 3a). The Eilers-Peeters curve is unimodal and right-skewed: growth rate increases with light intensity at low light intensities, reaches a maximum and then falls due to photoinhibition. It is written as:

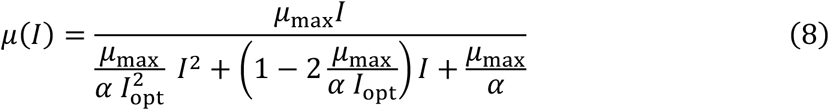

Here, *I* denotes the light intensity, *α* denotes the slope of the curve at *I* = 0 and *I*_opt_ denotes the optimum light intensity where growth rate *μ* = *μ*_max_.

The Norberg function is a left-skewed unimodal curve that describes per-capita growth rate (*μ*(*T*)) of ectotherms against temperature, *T* (Fig. 4a):

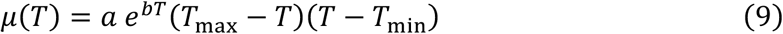

Here, *a* is the baseline Eppley growth rate (*a e^bT^*) at *T* = 0°C, *b* controls the rate of increase of the curve at low temperatures and *T*_min_ and *T*_max_ represent the minimum and maximum temperature at which the growth rate is zero.

The log-logistic function is a dose-response curve that describes how the per-capita growth rate (*μ*(*C*)) of a species changes with toxin concentration, *C* (Fig. 5a). This is an S-shaped, monotonically decreasing curve. Growth rate is maximum at *C* = 0 and as *C* → ∞, *μ*(*C*) → 0. We ignored hormesis wherein a small benefit is observed at low toxin concentrations. We used the three-parameter log-logistic model:

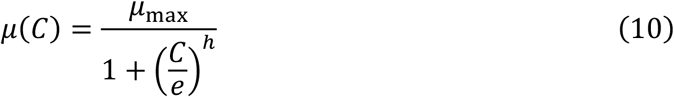

The three parameters are: maximum growth rate *μ*_max_, the effective concentration *e* where the growth rate is half of the maximum (commonly called EC50) and the slope ℎ of the curve at *C* = *e*.

All parameters and variables are summarized in Table 2. The prior distributions used for all parameters are described and plotted in Section 3, SI.

**Table 1:** A glossary of terms as they are used in the study.

| Term | Explanation |
| --- | --- |
| Categorical design | Design intended to be analyzed with an ANOVA, typically two levels |
| Gradient design | Design intended to be analyzed with a regression (linear or nonlinear), typically two levels |
| Driver | The manipulated (independent) variables: nutrients (or food), light, temperature and toxins. |
| Response | The measured quantity of interest: population growth rate or performance. |
| Experimental units | The smallest independently treated entity in an experiment e.g., a single chemostat. |
| Levels | Values of a driver in the experiment (e.g., 10°C). |
| Replicates | The number of experimental units at a level |
| Design space | The set of all possible levels. Our design space is a discrete grid of driver values. |
| Scoring rule | A summary quality measure that evaluates a predictive distribution. |
| Strictly proper scoring rule | A scoring rule whose expectation is minimized *only* when the predictive |
|  | distribution matches the distribution of the true value. |

**Table 2:** Units and definitions of parameters and statistical quantities.

| Symbol | Unit | Driver | Meaning |
| --- | --- | --- | --- |
| $S$ | $\mu\text{M}$ | Nutrient | Nutrient concentration |
| $I$ | $\mu\text{mol photons m}^{-2} \text{ s}^{-1}$ | Light | Light intensity<br>(Irradiance) |
| $T$ | $^{\circ}\text{C}$ | Temperature | Temperature |
| $C$ | $\mu\text{G L}^{-1}$ | Toxin | Toxin concentration |
| $\mu$ | $\text{day}^{-1}$ | All four | Per-capita growth rate |
| $\mu_{\max}$ | $\text{day}^{-1}$ | Nutrient, Light<br>and Toxin | Maximum per-capita<br>growth rate |
| $K$ | $\mu\text{M}$ | Nutrient | Half-saturation<br>constant; nutrient<br>concentration where<br>$\mu = \mu_{\max}/2$. |
| $I_{\text{opt}}$ | $\mu\text{E m}^{-2} \text{ s}^{-1}$ | Light | Optimum light intensity<br>i.e. $I$ where $\mu = \mu_{\max}$. |
| $\alpha$ | $\text{day}^{-1} (\mu\text{E m}^{-2} \text{ s}^{-1})^{-1}$ | Light | Slope at $I = 0$. |
| $a$ | $\text{day}^{-1} ^{\circ}\text{C}^{-2}$ | Temperature | Eppley growth rate<br>($a e^{bT}$) at $T = 0^{\circ}\text{C}$ |
| $b$ | $^{\circ}\text{C}^{-1}$ | Temperature | Rate of increase of Norberg curve at low temperatures |
| $T_{\min}$ | $^{\circ}\text{C}$ | Temperature | Temperature below which growth rate becomes negative |
| $T_{\max}$ | $^{\circ}\text{C}$ | Temperature | Temperature above which growth rate becomes negative |
| $e$ | $\mu\text{G L}^{-1}$ | Toxin | Effective concentration (EC50) where $\mu = \mu_{\max}/2$. |
| $h$ | Dimensionless | Toxin | Slope of the log-logistic function at $x = e$. |
| $g$ | $\text{day}^{-1}$ | | Function relating a driver to the conditional mean growth rate. |
| $x_i$ | Units of the focal driver | | $i$-th experimental unit. |
| $\epsilon_i$ | $\text{day}^{-1}$ | | Error of $i$-th growth rate measurement |
| $\sigma$ | $\text{day}^{-1}$ | | Standard deviation of the error distribution. |
| $n$ | | | Number of experimental units in a design. |
| $\theta$ | | | Parameter vector to be estimated. Does not include $\sigma$. |
| $p$ | | | Number of parameters in $\theta$. |
| $\psi^T$ | | | Vector of all parameters including $\sigma$. |
| $\delta$ | | | A design; a set of all experimental units $(x_1 \dots x_n)^T$ |
| $X$ | | | The design matrix. |
| $I$ | | | Fisher information matrix. |
| $u_{\text{SIG}}$ | nat | | The Shannon Information Gain (SIG) between a prior and a posterior. |
| $U_{\text{SIG}}$ | nat | | The expectation of the SIG value over the joint distribution of the parameters and the future data. |
| *CRPS* | Units of the focal parameter |  | Continuous Ranked Probability Score, a strictly proper scoring rule used to assess a marginal posterior. |
| *ES* | Dimensionless |  | Energy Score, an extension of CRPS to joint posterior distributions. |

### 2.4 SIG-optimal design calculation

For each model, we calculate SIG-optimal designs with 5, 7, 10 and 15 experimental units. We define a design space for each function that matches typical experimental practice for temperate and subtropical algae. For the nutrient, light, and temperature curves, the design space was an evenly spaced grid. For the toxin design space, we appended 0 to a log-scale grid between 0.1 and 1000.

We start the ACE algorithm with 40 randomly generated designs that run in parallel. At the end, we pick the design with the highest expected SIG utility (*U_SIG_*). The settings for the ACE algorithm are provided in Section 2, SI. The scripts are provided in the associated Github repository. We also plot the sensitivities (Eq. 3; Figs. S9-12) of all four functions with respect to their parameters to gain intuition about the placement of points in an optimal design.

### 2.5 Design evaluation

To evaluate the performance of the optimal designs, we compare them against uniform designs that are standard in these experiments. For every sample size, we compare the optimal design against a uniform design with the same number of experimental units. The uniform designs consisted of equally spaced experimental units across the design space without any replication for the Monod, Eilers-Peeters and Norberg functions. For a *n*-point uniform dose-response design, we ranked all points in the design space and chose *n* equally spaced points (in terms of rank) from between them; all uniform designs therefore included both 0 and 1000.

We compare the two designs on two fronts: parameter estimation and predictive accuracy. A SIG-optimal design is optimized to maximize information gain in the posterior. We test if this translates to better posteriors using scoring rules (Gneiting & Raftery, 2007) that summarize both the calibration and the precision of the posteriors. Finally, we test the optimal designs on their predictive ability as well, even though SIG-optimality does not optimize for predictions.

#### 2.5.1 Simulation experiments for design evaluation

For each function, we randomly drew 1000 parameter sets from the prior distributions and visually inspected the corresponding curves to ascertain that they were realistic (Figs. S2, 4, 6 and 8). For each combination of function, design type, sample size, and parameter set, we generated a simulated dataset assuming normally distributed errors (4 functions x 2 design types x 4 sample sizes x 1000 parameter sets = 32,000 simulated datasets). We estimated the function parameters in every simulated dataset with a Bayesian fitting approach using the R package *brms* (Bürkner, 2017). Details of the fitting procedure are provided in Section 4, SI.

**Figure 6:**
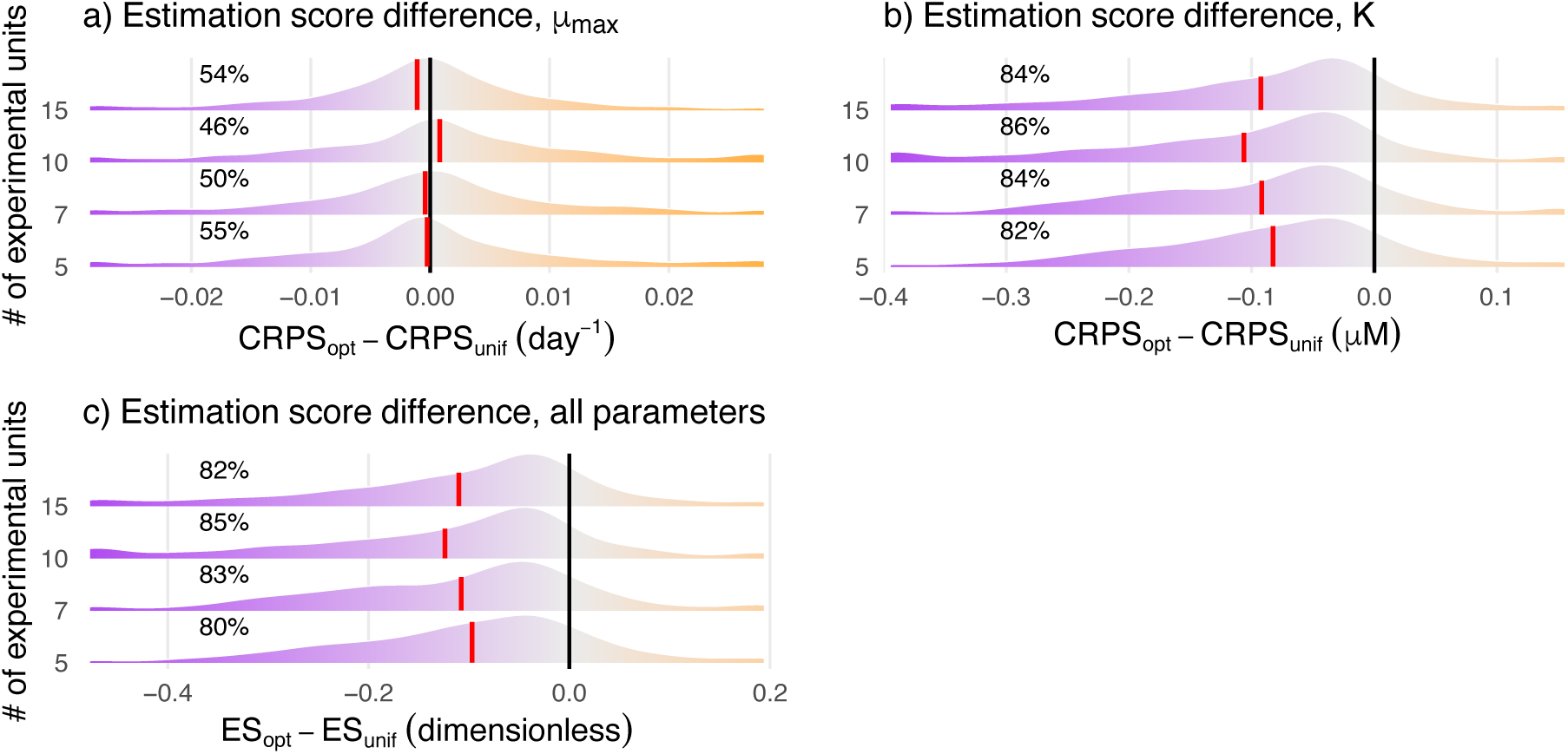
Ridgeline plots showing the distribution of differences in the CRPS and ES of the optimal and uniform designs for the Monod function with bad priors, across 1000 paired simulations at each sample size. The distributions have been trimmed such that values in the bottom and top 2 percentiles of all 4000 samples are pooled together at the edge of the plots. The black bar shows a difference of zero and the red bar denotes the mean difference between the scores. Negative values (to the left of the black bar) indicate simulations where the optimal design performed better and are shown in purple. The percentage of simulations where optimal design performs better is shown on the left of the plots. Even with bad priors, optimal designs are consistently better for estimating K and joint estimation of all parameters. Optimal designs are slightly better for estimating μ_max_.

#### 2.5.2 Evaluating parameter estimation error using proper scoring rules

For each simulation and model fit, we compare the posteriors resulting from uniform and optimal designs using a *strictly proper scoring rule* (Gneiting & Raftery, 2007). Although parameter estimation error is often compared using metrics such as mean absolute error (MAE), such metrics rely on point estimates and neglect the full posterior distributions from Bayesian fitting.

Posteriors can be ‘bad’ by either being inaccurate (mean/median is far from true value) or by being imprecise (extremely wide). A scoring rule is a summary measure that penalizes both. In our context, a scoring rule asks: how good was the posterior *π*(***θ***|***μ***, ***δ***) from a design ***δ*** given the true values ***θ***_true_ used for a simulation? Specifically, we compare the marginal posterior of a parameter *θ_i_* against the true value for that parameter *θ_i_*_,true_ using the *Continuous Ranked Probability Score* (CRPS) (Hersbach, 2000; Matheson C Winkler, 1976).

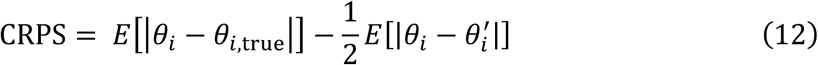

The first term is the Mean Absolute Error (MAE) that measures how far, on average, a draw from the posterior *θ_i_* is from the true value *θ_i_*_,true_. The second term is an adjustment term that rewards greater uncertainty when MAE is high. Lower CRPS values are better, rewarding posteriors that maximize sharpness while being well-calibrated (Gneiting C Raftery, 2007).

The CRPS is a generalization of the MAE to probabilistic distributions. If the posterior is deterministic (point value with 100% probability), the CRPS collapses to the MAE. Therefore, a probabilistic distribution with CRPS = *q* is equivalent in quality to a deterministic posterior with MAE = *q*. Furthermore, the CRPS is at its lowest value of zero if and only if the MAE is zero (point value = true value). There is no upper limit on the CRPS and it has the same units as the parameter being examined.

While CRPS is defined only for single parameters, the *Energy Score* (ES) is a multivariate generalization of the CRPS and can be used to assess the joint posterior (Gneiting C Raftery, 2007).

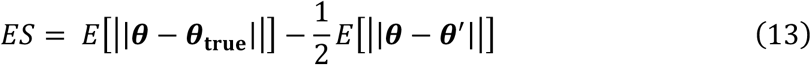

Here, ***θ*** and θ′ represent two draws from the joint posterior and ***θ*_true_** represents the vector of true values. Since calculating ES involves combining parameters on different scales, we standardized both the posterior samples and the true values and then calculated the energy scores using the standardized values. As with the CRPS, lower ES values indicate better joint posterior distributions. We calculate the CRPS and ES for both the uniform and optimal designs for each of the 1000 simulations for each function and sample size (complete distributions in Section 7, SI). We then compare the resulting distributions of CRPS and ES to assess if the optimal design outperforms the uniform design.

#### 2.5.3 Evaluating prediction error

Although SIG-optimal design is not designed to minimize prediction error, we chose to additionally compare prediction error across the two designs. For every simulation, we found a central estimated curve from the posterior distributions using the Modified Band Depth (MBD; Section 9, SI; López-Pintado & Romo, 2009). We compared the central estimated curve against the true curve by evaluating both curves’ prediction error at every point in the design space not contained in either of the two designs. We then calculated Mean Absolute Error (MAE) across the design space for each simulation for each design and compared the distribution of MAE values from each design. We refer to the MAE as the aggregate prediction error since it combines errors from points across the entire design space. We additionally calculated a point-wise MAE across all simulations i.e. the MAE across all simulations calculated separately for every evaluation point without averaging within a simulation (Figs. S18-21). This latter calculation allowed us to evaluate how prediction error varies across the driver based on the design.

### 2.6 Software

We used R version 4.6.0 (R Core Team, 2026) for the analyses. We also used ChatGPT versions o3 to 5.3 (OpenAI), Claude Opus 4.7 (Anthropic) and Gemini 3.1 Pro (Google) to assist in writing the code, but verified and modified the resulting code to ensure accuracy.

## 3 Results

Optimal designs differed strongly from uniform designs and sometimes involved a mix of replicated and unreplicated experimental levels. Optimal designs generally outperformed uniform designs in both parameter estimation and prediction error, especially for the two monotonic functions (nutrients and toxins).

### 3.1 Nutrients (Monod function)

SIG-optimal Monod designs (purple dots in Fig. 2b) allocate experimental units to two regions: around the half-saturation constant and at the maximum nutrient concentration in the design space, *S*_max_. The sensitivities of Monod function w.r.t *K* and *μ*_max_ are highest around *K* and *S*_max_ respectively as well (Fig. S9). Therefore, experimental units placed around the likely *K* and *S*_max_ strongly inform the inference of *K* and *μ*_max_ respectively. BOED accounts for the uncertainty in the parameters by spreading experimental units around the likely *K*. In contrast, *μ* = 0 at *S* = 0, so allocating experimental units at *S* = 0 provides no information at all.

The median CRPS across our 1000 simulations is lower for the optimal design for both *μ*_max_ and *K*. Optimal designs had a lower CRPS for *K* in more than 80% of the simulations at all sample sizes (Figs. 2c-d and S13b). In fact, the median CRPS for *K* of a 5-point optimal design is lower than the median CRPS of a 15-point uniform design (Fig. S13b). When the standardized joint posteriors are compared using the ES, the optimal designs again outperform uniform design in more than 80% of the simulations (Figs. 2e, S13c).

Median aggregate prediction error is lower for the SIG-optimal design across sample sizes (Figs. 2f). The point-wise prediction error follows the placement of units in the designs. Therefore, the point-wise prediction error is much lower for the SIG-optimal design at low nutrient levels but is similar to that of uniform design at medium-to-high levels (Fig. S18).

### 3.2 Light (Eilers-Peeters function)

SIG-optimal Eilers-Peeters designs have points in three broad regions: near the lowest light level, in the low-to-mid range around the *I*_opt_ prior, and near the highest light intensity value in the design space (Fig. 3b). The Eilers-Peeters function is most sensitive to changes in *α* at low light intensities (Fig. S10), so experimental units at low light intensities strongly inform the estimation of *α*. The sensitivity of the function is also high w.r.t *I*_opt_ and *α* near the highest light intensities (Fig. S10). Measurements near the highest light intensities thus inform the estimation of *α* and *I*_opt_. Finally, the magnitude of growth rate sensitivity w.r.t *μ*_max_is highest at *I*_opt_ and w.r.t *I*_opt_ has an optimum at a fraction of *I*_opt_ (Fig. S10). So, the remaining experimental units between the low light intensities and the likely *I*_opt_ contribute strongly to the estimation of *μ*_max_ and *I*_opt_. As the number of experimental units increase, the optimal design adds more points into these three regions instead of placing them elsewhere (Fig. 3b). Like the Monod function, measurements at *I* = 0 don’t provide any information since growth is always zero there.

Due to the allocation of more experimental units at low and the highest light intensities in the optimal design, the optimal designs outperform the uniform designs in the estimation of *α* and *I*_opt_, with lower median CRPS for both parameters (Figs. 3d-e and S14b-c). As the number of experimental units in the design increases from 5 to 15, optimal design adds more units at extremes while the uniform design ‘wastes’ more experimental units in low-sensitivity intermediate regions. Thus, the difference between the designs in CRPS for *α* and *I*_opt_ increases (Figs. 3d-e). In fact, the median CRPS for *α* from the optimal 5-unit design is lower than that from the uniform 15-unit design (Fig. S14b). The low-to-mid light intensities are captured reasonably well by a uniform design, so the differences in CRPS of *μ*_max_ are small (Figs. 3c and S14a). Finally, the optimal design ES is lower than that of the uniform design, indicating that its estimation of the standardized joint posterior is superior (Fig. 3f and S14d).

Unlike the Monod function, aggregate prediction error is similar between designs (Fig. 3g, S14e). Following the placement of units in the designs, the point-wise prediction error is substantially lower for optimal design at low-to-medium light levels but is slightly higher at medium-to-high levels (Fig. S19).

### 3.3 Temperature (Norberg function)

The SIG-optimal designs assign points to three regions: at the two extremes of the design space and around the peak of the curve (Fig. 4b). Given our prior, the sensitivities of the Norberg curve w.r.t *T*_min_ and *T*_max_ are usually highest at the corresponding extremes (Fig. S11), so growth measurements at the extremes constrain the estimation of *T*_min_ and *T*_max_. The points around the location of the peak contribute primarily to the estimation of *a* and *b*. The Norberg function is also most sensitive w.r.t *a* and *b* either at the location of the peak for *a* or after it for *b* (Fig. S11).

The CRPS values of the optimal design are substantially lower than the uniform design for both *T*_max_ and *T*_min_ (Figs. 4e,f and S15c,d). Thus, the optimal design estimates the boundaries of the thermal niche better. The designs are similar for *b* and also for *a* at low numbers of experimental units, with a slight advantage for the uniform design in the case of *a* as experimental units increase (Figs. 4 c,d and S15 a,b). The ES is lower for the optimal design as well (Fig. 4g), indicating it provides better parameter estimates overall.

As with the Eilers-Peeters function, aggregate prediction error is similar between designs (Fig. 4h, S15f) but point-wise prediction error follows the designs. The point-wise prediction error is lower for optimal design near the temperature extremes, but slightly higher at intermediate temperatures (Fig. S20).

### 3.4 Toxins (Log-logistic function)

All SIG-optimal designs allocate some experimental units to a toxin concentration of 0 to constrain the value of *μ*_max_ (Fig. 5b). The sensitivity of the log-logistic function w.r.t *μ*_max_ is also highest at a toxin concentration of 0 (Fig. S12). Since growth rate always goes to zero at infinitely high toxin doses, allocating units to high doses does not provide much information for most curves drawn from the prior. Instead, the optimal design aims to gain information about *e* and ℎ by placing units at low-to-intermediate concentrations. This is corroborated by the sensitivity optima of the log-logistic function w.r.t *e* and ℎ in the low-to-intermediate concentrations (Fig. S12). As the number of units increase, the optimal design adds points at higher doses to cover more unlikely scenarios based on the priors (Fig. 5b).

The median CRPS is lower for optimal design for EC50 and slope parameters (*e* and ℎ) (Figs. 5d-e, S16b-c), with the highest advantage at low numbers of experimental units. Interestingly, uniform designs have a lower median CRPS for *μ*_max_ (Figs. 5c, S16a). Optimal designs assign more experimental units to the low-to-intermediate regions (between 10 and 100) than the uniform designs, which assign more units at very low concentrations due to being evenly spaced at the logarithmic scale. Therefore, the optimal design trades off estimation of *μ*_max_ for better estimation of *e* and *h*. Optimal designs also have a lower median ES, indicating better parameter estimation overall with the strongest advantage at low numbers of experimental units (Fig. 5f, S16d).

As with the Monod function, median aggregate prediction error is lower for the optimal design (Fig. 5g). However, the difference between designs decreases substantially with sample size. This is borne out by the point-wise prediction error that follows the placement of units in the designs (Fig. S21). Optimal design is very slightly worse at both low and high toxin concentrations while being substantially-to-moderately better at intermediate concentrations based on sample size.

### 3.5 Bad priors

Our simulations thus far have relied on perfect priors: the true values for the simulation are drawn from the prior distributions used in fitting. This is unlikely to occur in reality, with many sources of bias causing scientists to develop priors that are at least moderately different from the truth. This leaves open the question: how well do optimal designs perform in more realistic conditions? To answer this, we repeated our simulations for the Monod designs but with true values being drawn from a distribution that was different from the priors used for fitting. In these new simulations, the true values of both *μ*_max_ and *K* were drawn from a lognormal distribution with arithmetic mean values 30% lower than that used for the priors (the standard deviation was unchanged). Across 1000 simulations, we found that the optimal design still outperformed the uniform design (Figs. 6, S17): CRPS for *K* was substantially lower (Figs. 6b, S17b) and for *μ*_max_ was similar between the two designs (Figs. 6a, S17a). The ES was again better for optimal design (Fig. 6c, S17c).

## 4 Discussion

Most experiments in ecology are designed by following Bayesian principles at least implicitly. Prior knowledge is often used to choose the question, the focal species, the specific driver and response and the design space. Experimentalists often know the function describing the relationship between the driver and response and understand intuitively that not all experimental measurements are equally informative. Bayesian experimental design formalizes this intuition and provides a framework to calculate designs that result in better posteriors on average (Chaloner & Verdinelli, 1995; Rainforth et al., 2024). BOED also circumvents the classic Catch-22 of optimal designs for non-linear regressions: calculation of optimal designs requires knowledge of parameters that cannot be known without performing the experiment. In this study, we applied BOED to four common single-driver growth experiments in ecology. All four are modelled as non-linear regressions that capture how growth (or performance) changes with nutrients, light, temperature and toxin levels. For each model, we calculated SIG-optimal designs with 5, 7, 10 and 15 experimental units.

Even for readers who do not wish to apply the full BOED procedure we described, our analyses offer some useful rules-of-thumb. Maximizing information gain in the parameters often translates to placing experimental units in regions of the design space where the curve is most sensitive to changes in parameters. For the unimodal functions (Norberg and Eilers-Peeters), allocating experimental units to the extremes and around the peak was particularly important to constrain parameters that control the rise, peak and fall of the growth curve. For the monotonic functions (Monod and log-logistic), the optimal designs constrained the asymptote by assigning points to the extreme with the maximum growth rate. The optimal designs also assigned experimental units to the regions where the curves change sharply: near the likely half-saturation constant for the Monod curve and the EC50 for the log-logistic curve. For all functions, prediction error is lowest in the regions where experimental units are allocated. These rules-of-thumb should help experimentalists develop custom, efficient designs that improve on existing practice even without applying BOED.

The optimal designs outperformed commonly used uniform designs on two fronts: 1) parameter estimation, which they are explicitly optimized for and 2) predictive accuracy, which they are not. For parameter estimation, we compared the performance of the two designs on the marginal posteriors (posteriors of individual parameters) and the standardized joint posteriors (of all parameters together). For all four models, the SIG-optimal design resulted in superior joint posteriors. It also outperformed the uniform design in the marginal posteriors for most parameters across the four functions. The two parameters where the uniform design had better estimates were *a* in the Norberg function and *μ*_max_ in the log-logistic function. In both cases, the optimal design trades off ability to estimate the focal parameter to improve estimation of the remaining parameters.

While SIG-optimal design is not focused on prediction error, this is important to consider because experimental parameter estimates are often implemented in dynamic models where prediction error matters. We found that aggregate prediction error was substantially lower for optimal design in the two monotonic functions – nutrients and toxins. For the two unimodal functions, it was similar between optimal and uniform designs. However, aggregate prediction error masked large underlying differences across the environmental gradients. The two designs offered lower prediction error in different regions of the environmental gradient corresponding to where they placed their experimental units. Therefore, when designing an experiment, it would be worthwhile to consider whether there are parts of the design space that are more important to predict accurately.

BOED can appear daunting to experimental biologists with limited mathematical training. To address this, we have provided detailed R code implementing the calculation and evaluation of SIG-optimal designs for the four models discussed here. The code will allow an interested experimenter to change the prior distributions, impose additional constraints on the design space and try out other models.

Despite the challenges, we believe that BOED offers large benefits. Perhaps the most important advantage of optimal designs is that they are more efficient (Thomas C Ranjan, 2024). They can offer the same statistical performance as a uniform design with fewer points and therefore reduce experimental effort. For some functions such as Monod, a SIG-optimal design with 5 experimental units performs as well as a 15-unit uniform design. A researcher could therefore take advantage of this efficiency gain to reduce their time and resource investment, or to expand the scope of their experiment – for example, to measure three species’ Monod curves instead of one. BOED will also pair well with the advent of robotics-driven automated high-throughput experiments, to measure important traits such as the half-saturation constant, the EC50 or the *T*_max_ for many species quickly and efficiently. Using these designs will allow the experimenter to conduct more advanced experiments that rely on response curves, such as experimental evolution and pairwise competition experiments.

Bayesian experimental design has never been used in ecological experiments to our knowledge. Our study is a first step in this direction, and numerous opportunities exist to further develop BOED methods for ecology experiments. Optimal designs are function-dependent and often multiple functions are used to describe growth response to the same driver. It would be useful to compare optimal designs across multiple such functions to see how much they change. We have also assumed that the errors are homoscedastic. Relaxing this assumption may result in a different optimal design (Song & Kee Wong, 1998). And while we focused on parameter estimation, designs can be optimized for prediction and model discrimination as well. On the computational side, exciting new machine learning algorithms can speed up the design calculation further (Rainforth et al., 2024). Overall, we believe that BOED presents an immense opportunity to speed up common experiments in ecology and make them more cost-effective.

## Acknowledgements

RR acknowledges support from the HIFMB Postdoc Program (HIPP) at HIFMB, a collaboration between the Alfred-Wegener-Institute, Helmholtz-Center for Polar and Marine Research and the Carl-von-Ossietzky University Oldenburg, initially funded by the Ministry of Science and Culture of Lower Saxony (MWK) and the Volkswagen Foundation through the ‘Niedersächsisches Vorab’ grant programme (ZN3285).

## Conflicts of Interest statement

The authors have no conflicts of interest to declare.

## Author contribution statement

Both authors conceptualized the study. RR wrote the code to calculate the designs, made Fig. 1 and wrote the first draft. MKT wrote the code to evaluate the designs and made the rest of the figures. Both authors interpreted results and wrote the revisions.

## Data availability

Code to reproduce the results is available at: https://github.com/raviranjan545/boed-paper-reproduction. Additional code to calculate and explore SIG-optimal designs is available at: https://github.com/raviranjan545/boed-calculation

## Statement of inclusion

Our study did not involve data, therefore no stakeholders were involved.

## Notes

### Competing Interest Statement

The authors have declared no competing interest.

### Summary of Updates

We fixed a few minor errors, updated the plots and the text.

