## Supplementary Information for "Improving the efficiency of single-driver ecological experiments with Bayesian optimal experimental design"

#### 1. Numerical evaluation of $U_{\mathrm{SIG}}(\boldsymbol{\delta})$

As shown in eq. 8 in the main text, the expected SIG utility $U_{\mathrm{SIG}}(\boldsymbol{\delta})$ of a design $\boldsymbol{\delta}$ is:

$$U_{\mathrm{SIG}}(\boldsymbol{\delta}) = \iint_{\boldsymbol{\theta},\boldsymbol{\mu}} u_{\mathrm{SIG}}(\boldsymbol{\theta},\boldsymbol{\mu},\boldsymbol{\delta})\, \pi(\boldsymbol{\mu}|\boldsymbol{\theta},\boldsymbol{\delta})\, \pi(\boldsymbol{\theta})\, d\boldsymbol{\theta}\, d\boldsymbol{\mu} \tag{S1}$$

Since the integral is nested, numerical evaluation of the expected SIG utility for a given design involves two nested loops. We provide a broad overview of the numerical procedure here. For a more detailed description, see Section 3.1of Overstall & Woods, 2017.

1. In the outer loop, a parameter set is drawn from the prior distribution. Using the parameter set and the likelihood, an artificial dataset is calculated.
2. Next, the inner loop is initiated to calculate the marginal likelihood, which is the probability density of the focal dataset marginalized across the prior distribution. Thousands of new parameter sets are drawn from the prior and the likelihood of the dataset under each of these parameter sets is calculated and averaged to generate the marginal likelihood.
3. Using the marginal likelihood from Step 2 and the likelihood of the data given the parameter set from Step 1, eq 7 (main text) can be used to calculate the information gained (SIG value) for that specific dataset.
4. Steps 1 through 3 are repeated thousands of times to calculate the SIG value for thousands of artificial datasets drawn from distinct parameter sets drawn from the prior. The SIG values are then averaged to get the expected SIG utility for the focal design.

To find an optimal design, steps 1 through 4 must be repeated for many different candidate designs to calculate their expected SIG utility and the best design must be chosen.

#### 2. Approximate Coordinate Exchange (ACE) algorithm

ACE is a two-phase stochastic optimization algorithm designed to find Bayesian optimal designs. Here, we provide a brief overview of the algorithm. For a more detailed description, see Overstall & Woods, 2017. In Phase I, ACE starts with an initial design and focuses on one experimental unit ($x_i$) at a time while freezing the others. It then evaluates the expected utility ($U_{\mathrm{SIG}}$ in our case) at a small set of candidate experimental units to construct a Gaussian Process (GP) emulator. Intuitively, a Gaussian process is a distribution over functions – effectively, a collection of possible continuous functions. The small set of evaluated $U_{\mathrm{SIG}}$ values is used as a training set that narrows this distribution down to a set of functions that are consistent with the evaluated $U_{\mathrm{SIG}}$ values. The narrow set of functions form a GP emulator of the original $U_{\mathrm{SIG}}$ function and allow us to calculate an approximate $U_{\mathrm{SIG}}$ for any other collections of experimental units outside the training set. The advantage of using a GP emulator is that it is fast and computationally inexpensive when compared to the complex original $U_{\mathrm{SIG}}$ function. The GP emulator is then maximized to suggest a new experimental unit. Thus, the Phase I algorithm goes through the entire design one experimental unit at a time and identifies the best replacement in the design space for each experimental unit.

In Phase II, the ACE algorithm focuses on replication, an often-desired characteristic of experimental designs. It does so by swapping points — adding replicates of experimental units that increase $U_{\mathrm{SIG}}$ the most and then deleting experimental units that leaves behind the design with highest $U_{\mathrm{SIG}}$. Error could creep into this process through the Monte Carlo approximation of the expected utility and the GP emulator. Therefore, the algorithm uses probabilistic Bayesian tests of equality to accept or reject changes in the design.

The ACE algorithm is started multiple times from different Latin hypercube initial designs to avoid local optima and the design with the highest expected SIG value is chosen. In this study, we started the ACE algorithm from 40 starting designs with random assignment of points and chose the best final design among them. We used the Monte Carlo method (method = “MC” in the *pacenlm* function from the *acebayes* package) with the criterion set to Shannon Information Gain (criterion = “SIG”). For each starting design, we set number of Phase I iterations to 40 (N1 = 40 in the code) and Phase II iterations to 200 (N2 = 200 in the code). The sample size for the Bayesian tests of equality is set to 40000 and the sample size for the Monte Carlo integration is set to 2000 (B = c(40000, 2000)). Finally, we modified the C++ code associated with the *acebayes* package that calculates the SIG utility value to prevent numerical underflow and overflow. We implemented a log-sum-exp operation within the code to prevent this problem. Note that the design calculation process in *acebayes* is inherently stochastic and not reproducible despite setting seeds. Therefore, each design calculation starting with the same starting design and the same settings will produce a slightly different final design.

#### 3. Prior distributions

We defined lognormal priors for most parameters. Lognormal distributions always result in positive parameters and have long upper tails on a linear scale. They are more realistic than Gaussian for these and many other biological parameters. They also demonstrate that we do not need to oversimplify our priors to calculate optimal designs. Priors for all parameters of a function were also all uncorrelated, except in the case of the Norberg function. For computational ease, we rescaled all parameters to have a mean of 1, with the Norberg function being an exception here as well. For the Norberg function, we recentered the temperature values to improve numerical stability during Bayesian model fitting. The recentering process for the Norberg function is described in Section 5 while Bayesian model fitting details are in Section 4.

We describe prior distributions here before the rescaling. Where functions shared common parameters, we used identical priors. Thus, both maximum growth rate ($\mu_{\max}$; common in three functions) and error standard deviation ($\sigma$; common in all four) have a common prior across functions.

$$\log \mu_{\max} \sim \mathcal{N}(0.01, 0.4^2) \tag{S2}$$

$$\log \sigma \sim \mathcal{N}(-2.3, 0.1^2) \tag{S3}$$

Below, we describe and plot the priors for the rest of parameters for each function. The prior distribution for $\mu_{\max}$ is plotted wherever the parameter is present. While $\sigma$ is present for all functions, we skip plotting it for temperature and focus on plotting the correlations between the Norberg parameters.

For each function, we also check the priors to see if they result in realistic curves by plotting 1000 curves using 1000 parameter sets drawn from the respective prior distribution(s). These 1000 parameter sets also function as the true values in the simulations used to evaluate the optimal and uniform designs (Section 2.5, main text and Section 4, SI).

#### 3.1 Nutrients (Monod function)

For the Monod function, the half-saturation constant $K$ also follows a lognormal distribution.

$$\log K \sim \mathcal{N}(0.3, 0.5^2) \tag{S4}$$

The prior distributions for both $\mu_{\max}$ and $K$ are plotted in Fig. S1 below. We plotted Monod curves in Fig. S2 using 1000 parameter sets drawn from the prior distributions.

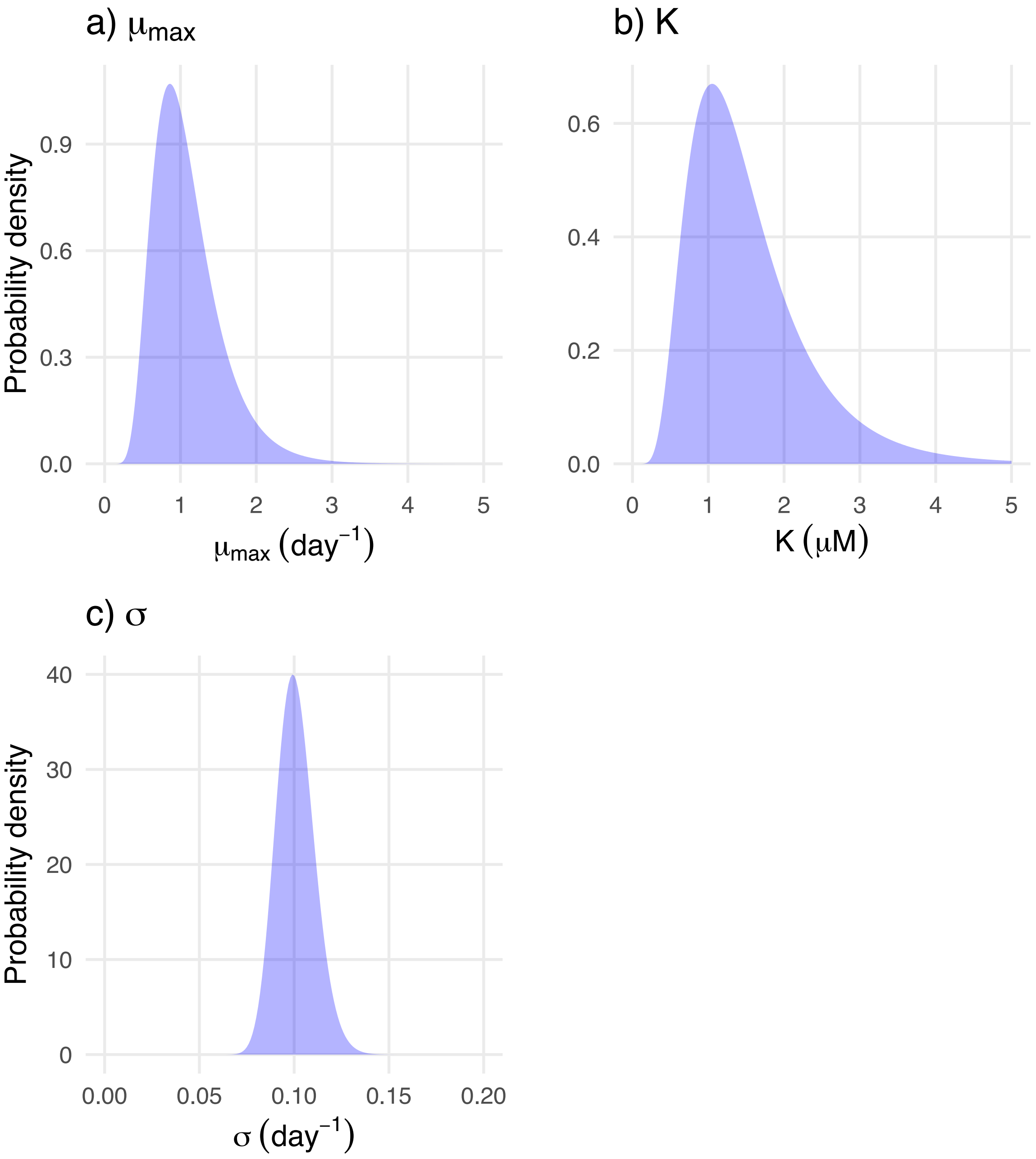

*Figure S1: Prior distributions for Monod parameters: a) Maximum growth rate $\mu_{\max}$ and b) half-saturation constant $K$.*

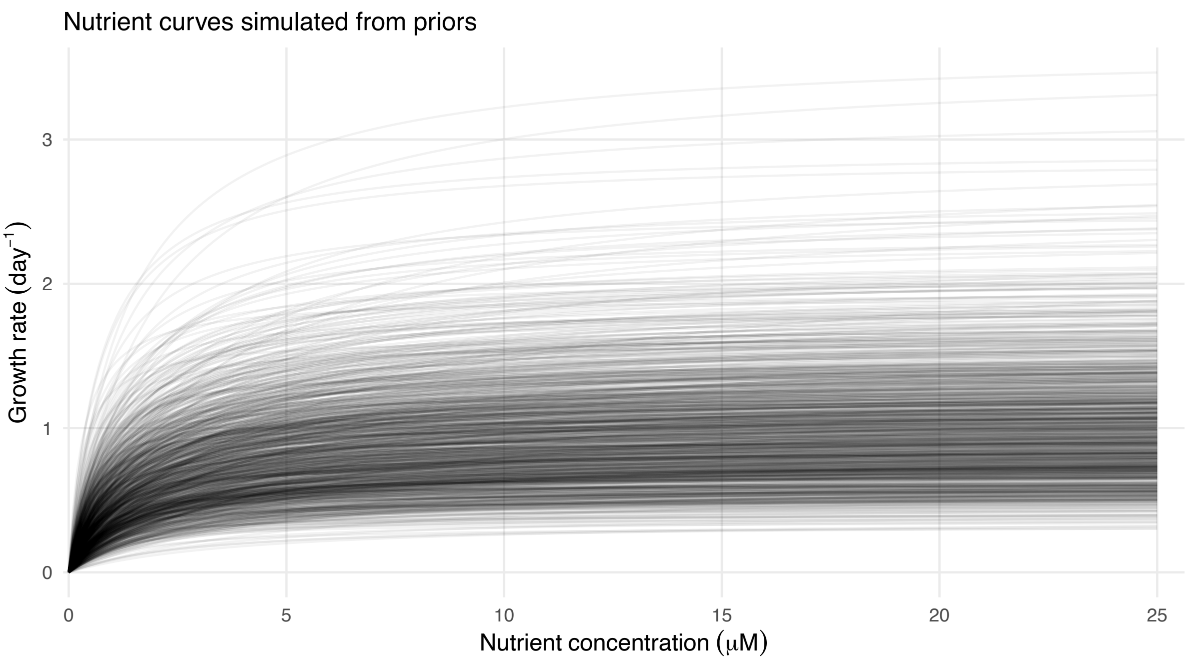

*Figure S2: Monod curves drawn using 1000 parameter sets drawn from the prior distributions.*

#### 3.2 Light (Eilers-Peeters function)

For Eilers-Peeters, $\alpha$ and $I_{\mathrm{opt}}$ follow lognormal distributions.

$$\log \alpha \sim \mathcal{N}(-3, 0.8^2) \tag{S5}$$

$$\log I_{\mathrm{opt}} \sim \mathcal{N}(5.5, 0.3^2) \tag{S6}$$

The prior distributions for $\mu_{\max}$, $\alpha$ and $I_{\mathrm{opt}}$ are plotted in Fig. S3 below. We plotted Eilers-Peeters curves in Fig. S4 using 1000 parameter sets drawn from the prior distributions.

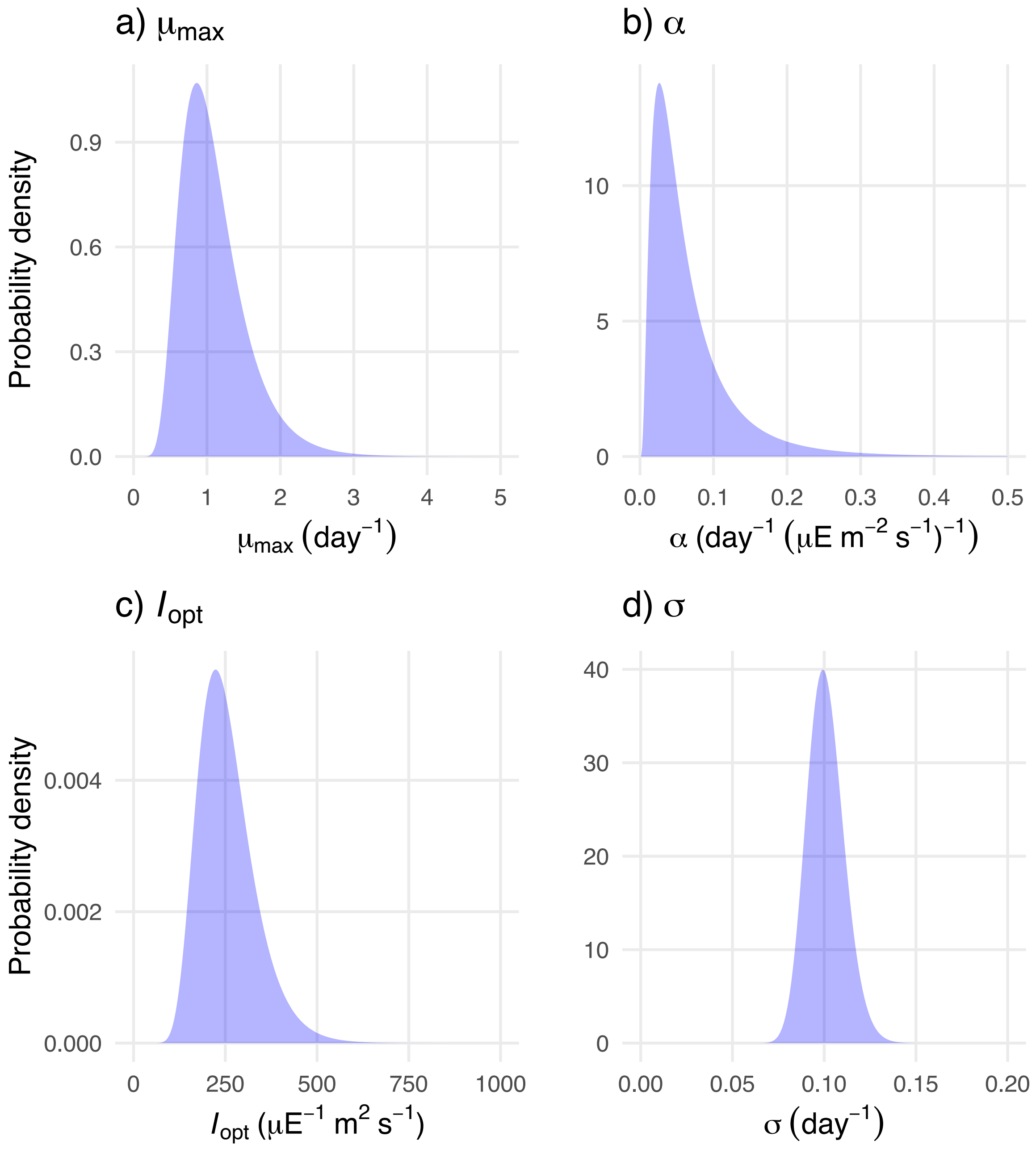

*Figure S3: Prior distributions for Eilers-Peeters parameters: a) Maximum growth rate $\mu_{\mathrm{max}}$, b) initial slope $\alpha$ and c) the optimum light intensity $I_{\mathrm{opt}}$.*

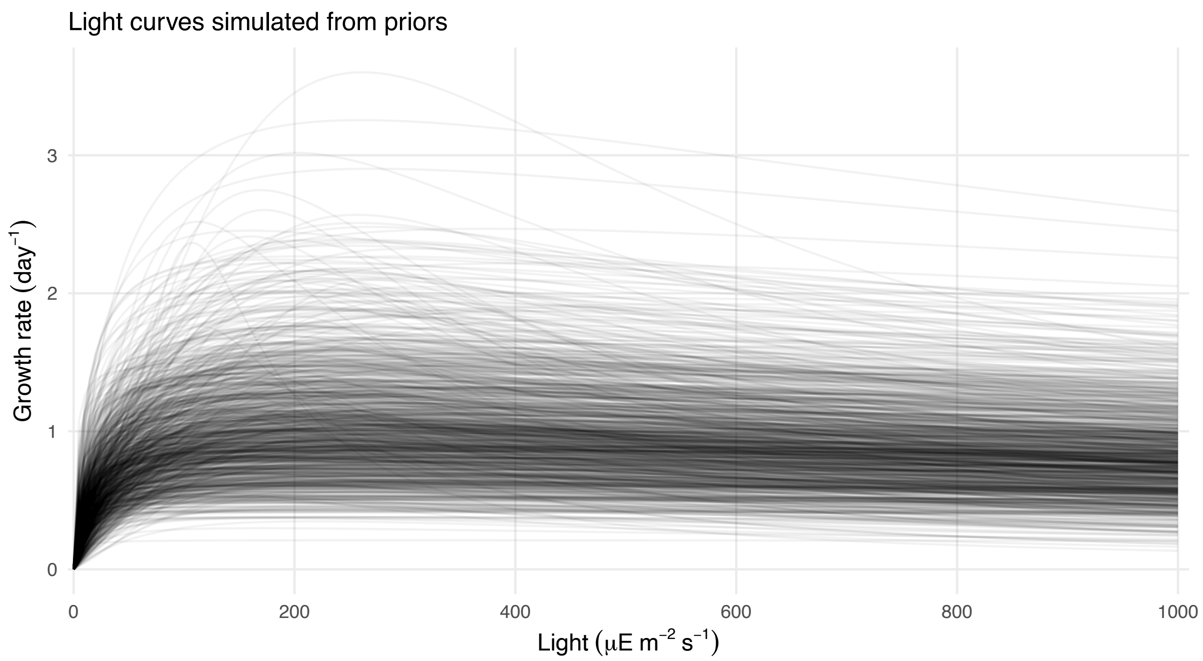

*Figure S4: Eilers-Peeters curves drawn using 1000 parameter sets drawn from the prior distributions.*

#### 3.3 Temperature (Norberg function)

Uncorrelated parameter values allow for temperature curves to take unrealistic shapes, with impossibly high maximum growth rates. We therefore defined a multivariate normal distribution as the prior for the Norberg function. We defined the marginal distributions for $\log a$, $\log b$, $T_{\min}$ and $T_{\max}$ as normal distributions and imposed a covariance matrix between the parameters to obtain a joint prior distribution that generates realistic temperature curves. Note that the marginals for $a$ and $b$ are lognormal while $T_{\min}$ and $T_{\max}$ are normal.

$$\boldsymbol{\theta}_{\text{Norberg}} \sim \mathcal{N}(\boldsymbol{m}, \mathbf{SCS}) \tag{S7}$$

Here, $\boldsymbol{\theta}_{\text{Norberg}}$ is the parameter vector and $\boldsymbol{m}$ is a vector with means of the marginal normal distributions. The matrix product $\mathbf{SCS}$ is a covariance matrix where $\boldsymbol{S}$ is a diagonal matrix with the standard deviations of the marginal normal distributions and $C$ is the correlation matrix.

$$\boldsymbol{\theta}_{\text{Norberg}} = \begin{pmatrix} \log a \\ \log b \\ T_{\max} \\ T_{\min} \end{pmatrix}, \boldsymbol{m} = \begin{pmatrix} m_{\mathrm{a}}^{*} \\ m_{\mathrm{b}}^{*} \\ m_{\mathrm{T_{max}}} \\ m_{\mathrm{T_{min}}} \end{pmatrix} \tag{S8}$$

$$\boldsymbol{S} = \operatorname{diag}\left(\sigma_{\mathrm{a}}^{*}, \sigma_{\mathrm{b}}^{*}, \sigma_{\mathrm{T_{max}}}^{*}, \sigma_{\mathrm{T_{min}}}^{*}\right) \tag{S9}$$

$$\boldsymbol{C} = \begin{pmatrix} 1 & -0.6 & -0.5 & 0.3 \\ -0.6 & 1 & 0 & 0 \\ -0.5 & 0 & 1 & 0 \\ 0.3 & 0 & 0 & 1 \end{pmatrix} \tag{S10}$$

For $a$ and $b$, ($m_{\mathrm{a}}^*$, $m_{\mathrm{b}}^*$) are the log-scale means and ($\sigma_{\mathrm{a}}^*$, $\sigma_{\mathrm{b}}^*$) are the log-scale standard deviations. These can be written in terms of the arithmetic scale means ($m_a$, $m_b$) and standard deviations ($\sigma_a$, $\sigma_b$) of a lognormal distribution:

$$\sigma_{\mathrm{a}}^* = \sqrt{\log\left(1 + \left(\frac{\sigma_{\mathrm{a}}}{m_{\mathrm{a}}}\right)^2\right)}, m_a^* = \log m_{\mathrm{a}} - \frac{1}{2}(\sigma_{\mathrm{a}}^*)^2 \tag{S11}$$

$$\sigma_{\mathrm{b}}^* = \sqrt{\log\left(1 + \left(\frac{\sigma_{\mathrm{b}}}{m_{\mathrm{b}}}\right)^2\right)}, m_b^* = \log m - \frac{1}{2}(\sigma_{\mathrm{b}}^*)^2 \tag{S12}$$

Finally, the arithmetic scale means and standard deviations are:

- $a$: $m_{\mathrm{a}} = 0.002, \sigma_{\mathrm{a}} = 0.0002$
- $b$: $m_{\mathrm{b}} = 0.025, \sigma_{\mathrm{b}} = 0.005$
- $T_{\mathrm{max}}$: $m_{\mathrm{T_{max}}} = 35, \sigma_{\mathrm{T_{max}}} = 2$
- $T_{\mathrm{min}}$: $m_{\mathrm{T_{min}}} = 35, \sigma_{\mathrm{T_{min}}} = 2$

The multivariate normal prior for Norberg parameters is shown in Fig. S5 below. We plotted Norberg curves in Fig. S6 using 1000 parameter sets drawn from the prior distributions.

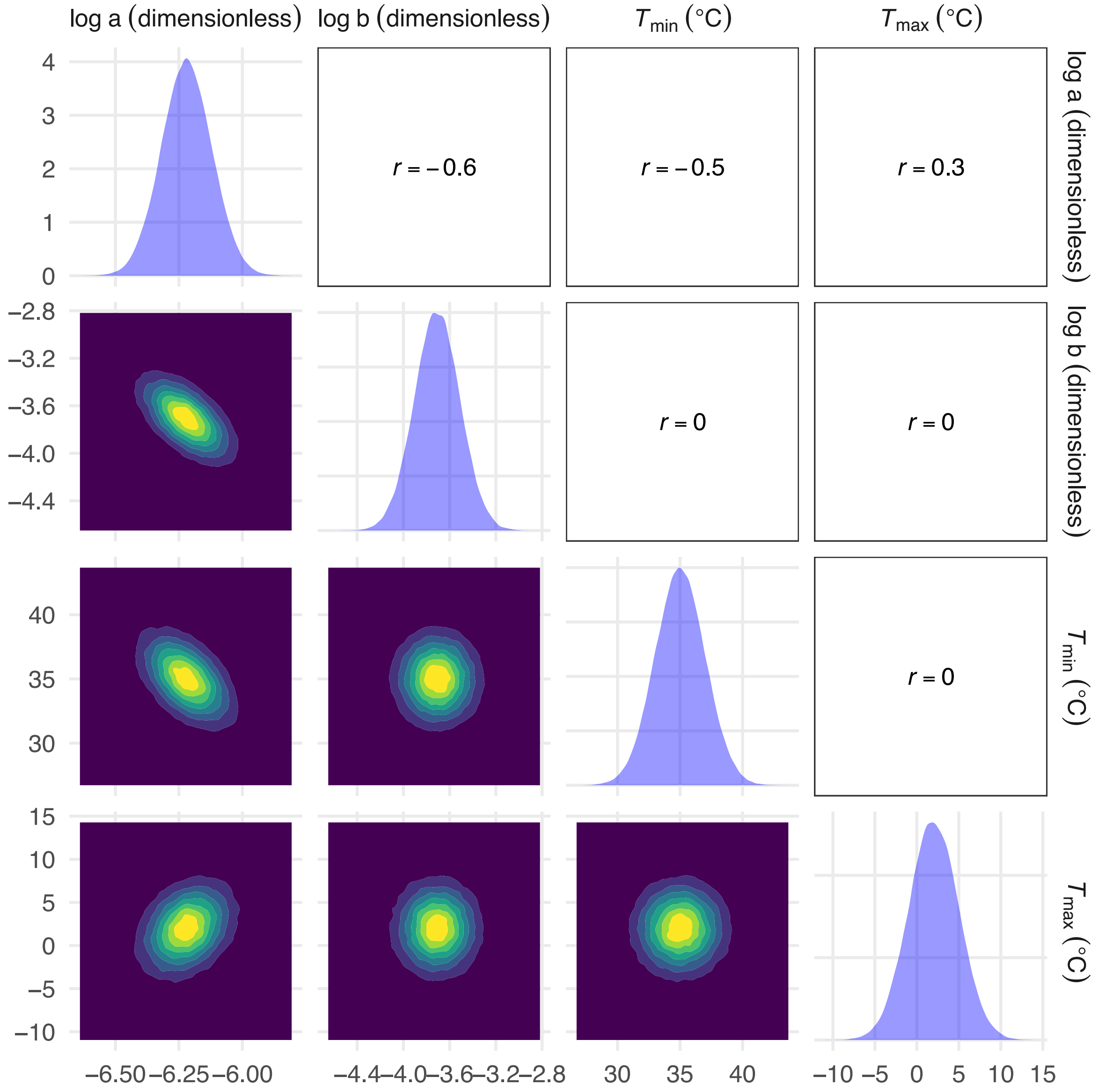

*Figure S5: The multi-variate normal prior for Norberg parameters. Plots on the diagonal show the marginal distributions of $\log a$, $\log b$, $T_{\min}$ and $T_{\max}$. The off-diagonal plots show the correlations imposed upon the focal pair of parameters.*

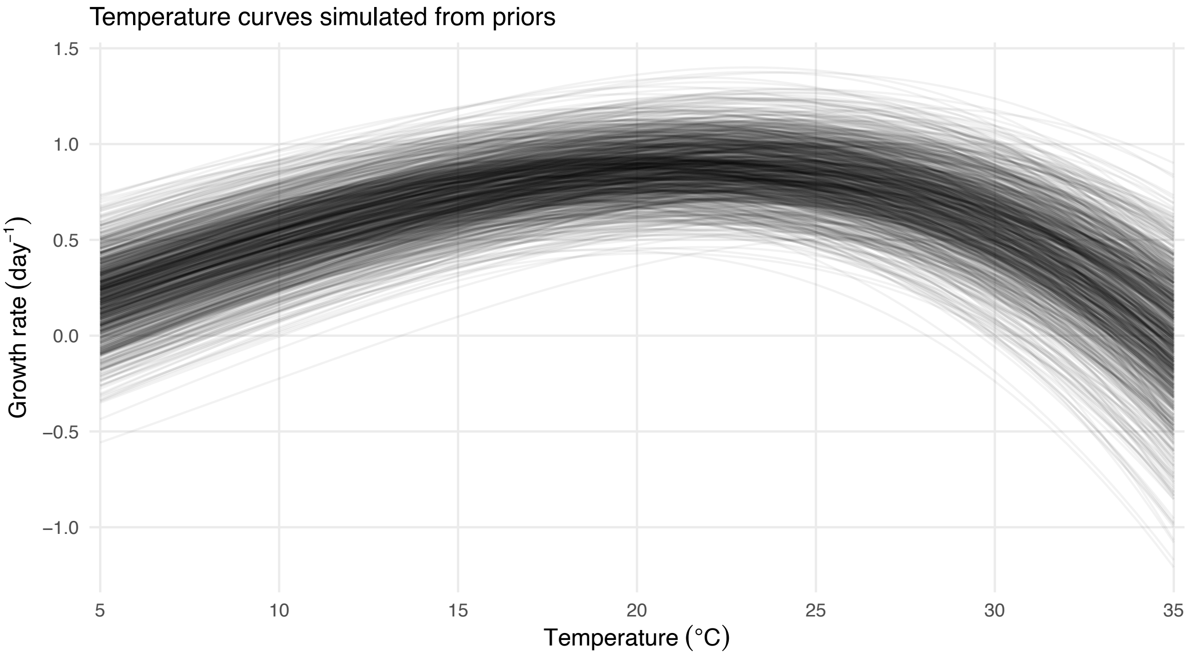

*Figure S6: Norberg curves drawn using 1000 parameter sets drawn from the multi-variate normal prior distribution.*

#### 3.4 Toxins (Log-logistic function)

For Eilers-Peeters, $e$ and $h$ follow lognormal distributions.

$$\log e \sim \mathcal{N}(4, 1^2) \tag{S13}$$

$$\log h \sim \mathcal{N}(1, 0.5^2) \tag{S14}$$

The prior distributions for $\mu_{\max}$, $e$ and $h$ are plotted in Fig. S7 below. We plotted log-logistic curves in Fig. S8 using 1000 parameter sets drawn from the prior distributions.

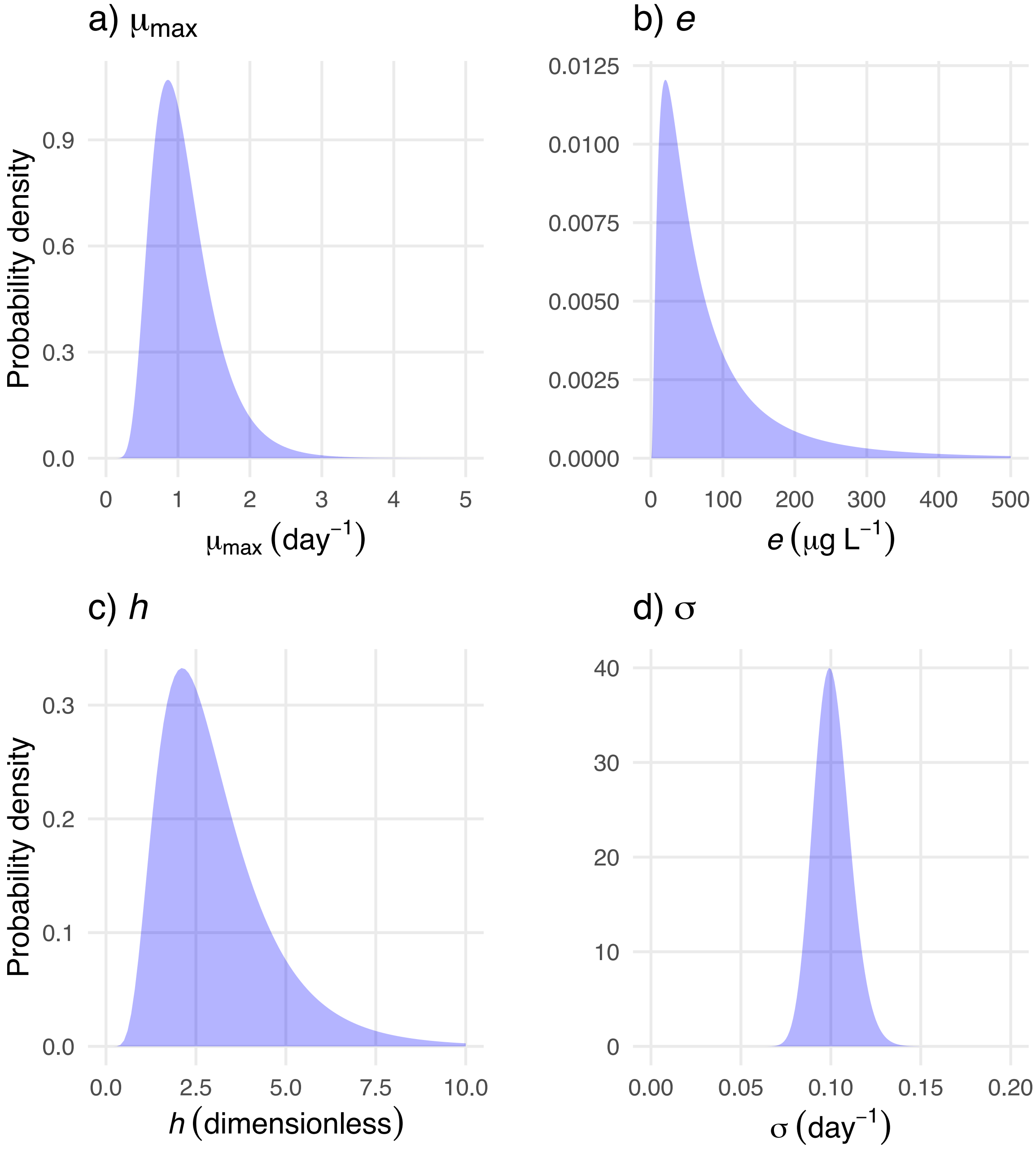

*Figure S7: Prior distributions for log-logistic parameters: a) Maximum growth rate $\mu_{\max}$, b) EC50, $e$ and c) slope parameter $h$.*

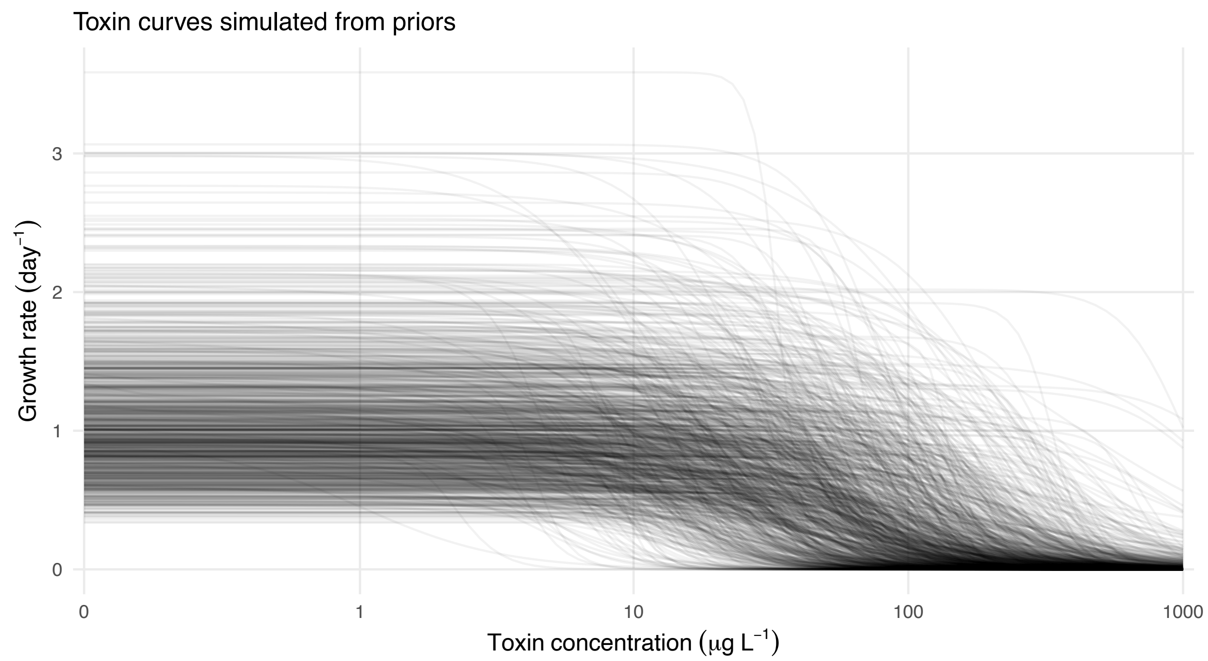

*Figure S8: Log-logistic curves drawn using 1000 parameter sets drawn from the prior distributions.*

#### 4. Bayesian parameter estimation from simulated datasets

We used the R package *brms* for Bayesian model fitting (Bürkner, 2017). Every fit used 4 chains with 5000 iterations each, half of which were discarded as warmup (or 'burn-in'). We specified different target average acceptance probabilities (0.95 – 0.99) and maximum tree depth (15 - 20) values for the different functions, as they varied in how easy they were to fit. We also provided the fitting functions with initial parameter guesses; we used the mean of each parameter's prior with a small amount of random noise added. We modified the Norberg function to ensure numerical stability during the Bayesian model fitting (details in Section 5 below).

We checked fit diagnostics for all fits: $\hat{R}$, bulk Effective Sample Size (ESS), tail ESS, and number of divergences. In a very small number of cases, there were a few divergences. We investigated these further and judged them to be unimportant based on $\hat{R}$ values, ESS values and visual examinations of the chains, posterior distributions and fitted curves.

#### 5. Recentering of temperature for Norberg analysis

In the Norberg equation, growth rate at low temperatures ($> T_{\min}$) is determined primarily by $\exp(\log a + b\,T)$. Since $a$ is the Eppley growth rate at $T = 0°C$ while the simulated experiment is conducted between 5°C and 35°C, slight changes in $b$ require large compensatory changes in $\log a$ to fit a given set of data points. Thus, $\log a$ and $b$ estimates have a strong negative correlation that can cause numerical instabilities in MCMC sampling during Bayesian model fitting. To counter this, we recentered temperature around a reference temperature $T_0$ during model fitting such that our new centered temperature variable $T' = T - T_0$. We rewrote the Norberg equation in terms of the recentered temperature:

$$\mu(T') = a\, e^{bT_0}\, e^{bT'} \left((T_{\max} - T_0) - T'\right) (T' - (T_{\min} - T_0)) \tag{S15}$$

The centering temperature $T_0 \approx 20.89°C$ is chosen as the mean of all the points from the uniform and optimal designs. We further define a joint prior distribution in terms of $\{\log a, \log b, T_{\max}, T_{\min}\}$ and estimate the posterior in these terms as well (Section 3.3, SI). This has the added advantage that only positive values of $a$ and $b$ are sampled by the MCMC algorithm. We plot the results after converting the estimates back to $\{a, b, T_{\max}, T_{\min}\}$.

#### 6. Sensitivities of all functions

Here, we plot the sensitivity of each function w.r.t each of its parameters. The sensitivity ($\partial g/\partial \theta_j$; Eq. 3, main text) of a function $g$ measures how much the predicted growth rate $g(x_i)$ measured at a level $x_i$ would change in response to changes in the focal parameter $\theta_j$. In case of the four functions used in this study, the sensitivities are themselves non-linear functions whose magnitude and sometimes even the shape depend on parameter values. We evaluated these sensitivities using the specific parameter set that generated the median functional curve. This representative curve was identified from prior predictive simulations using Modified Band Depth (MBD, Section 9). The specific parameters are listed in Table S1, Section 9. Figures S5-8 show the sensitivities of the Monod, Eilers-Peeters, Norberg and the log-logistic function.

The sensitivity curves plotted here are only meant to be representative. In case of the Norberg sensitivity to $T_{\min}$, the shape can change from monotonic to non-monotonic based on the parameter values chosen. The non-monotonic shape of the Norberg sensitivity to $T_{\min}$ occurs under parameter combinations that have a probability of about 0.17 under the prior we are using.

While we use these sensitivity plots to understand the placement of experimental units in the SIG-optimal design, it is important to note that an optimal design for a function is not simply a collection of levels corresponding to the sensitivity optima of its parameters. The parameters are often structurally correlated within the model, meaning that the Fisher Information Matrix (eq. 4, main text) contains off-diagonal elements that dictate trade-offs in where experimental units should be placed. Further, if the priors impose additional correlations among the parameters (such as in the Norberg function), that might also impact the optimal design. Therefore, the sensitivity plots are meant only to be used as a guide to identify important regions of the design space where an optimal design might place experimental units.

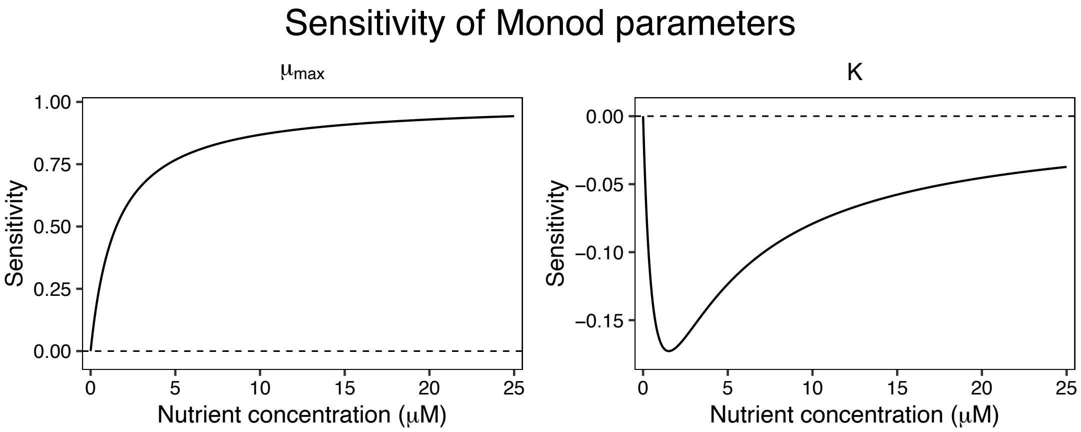

*Figure S9: Sensitivity of the Monod function to maximum growth rate $\mu_{max}$ and half-saturation constant $K$.*

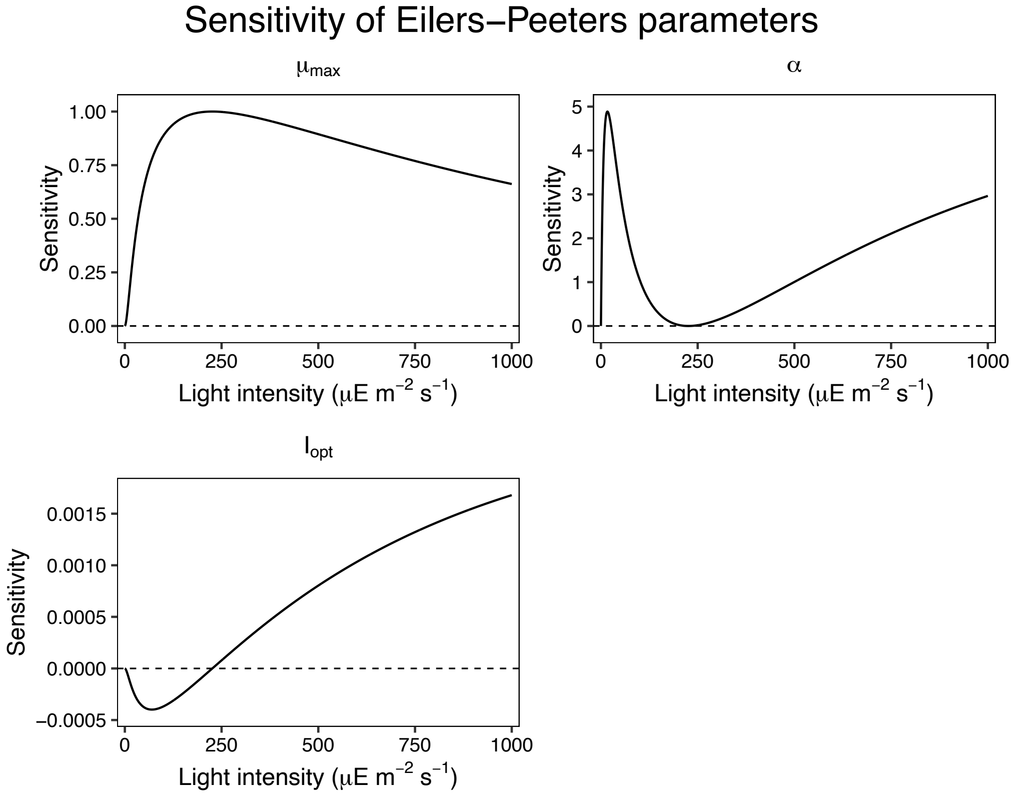

*Figure S10: Sensitivity of the Eilers-Peeters to maximum growth rate $\mu_{max}$, initial slope $\alpha$ and the optimum light intensity $I_{opt}$.*

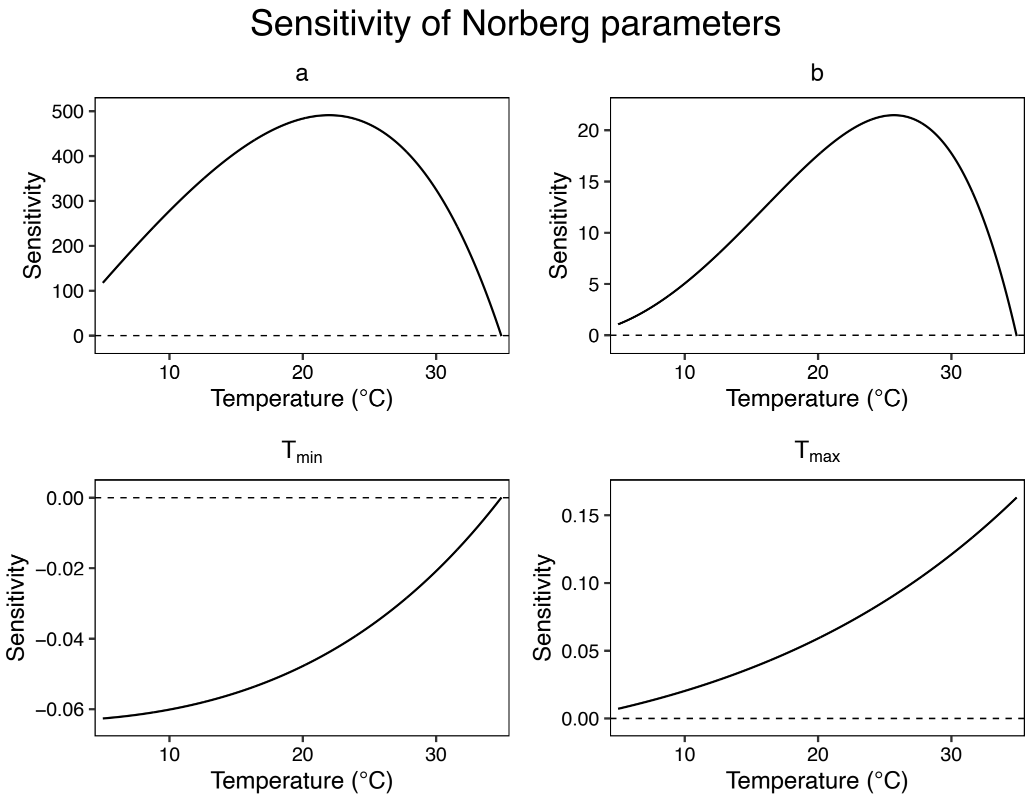

*Figure S11: Sensitivity of the Norberg function to baseline Eppley growth rate at T=0°C, a, the rate of increase of the curve at low temperatures, b, the maximum temperature at which the growth rate is zero,* $T_{max}$ *and the minimum temperature at which the growth rate is zero,* $T_{min}$*.*

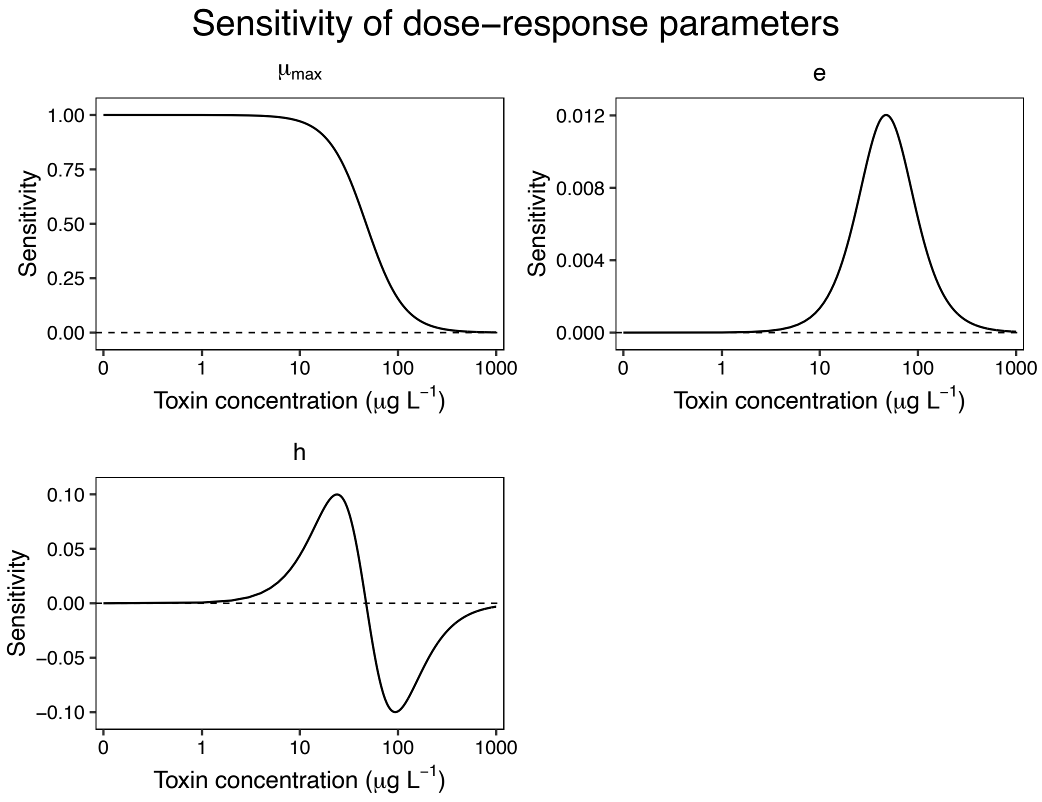

*Figure S12: Sensitivity of the log-logistic function to maximum growth rate $\mu_{max}$, the EC50 $e$ and the slope parameter $h$.*

### 7. Additional CRPS and ES plots

In the main text, we showed the difference in CRPS and ES for parameter estimation and aggregate prediction error between the optimal and uniform designs for all four functions (Figs. 2-6). Here, we provide the distributions of CRPS and energy scores for each function. Note that due to the long tails, we plot the scores on a log axis. Figures S9-12 show the sensitivities of the Monod, Eilers-Peeters, Norberg and the log-logistic function.

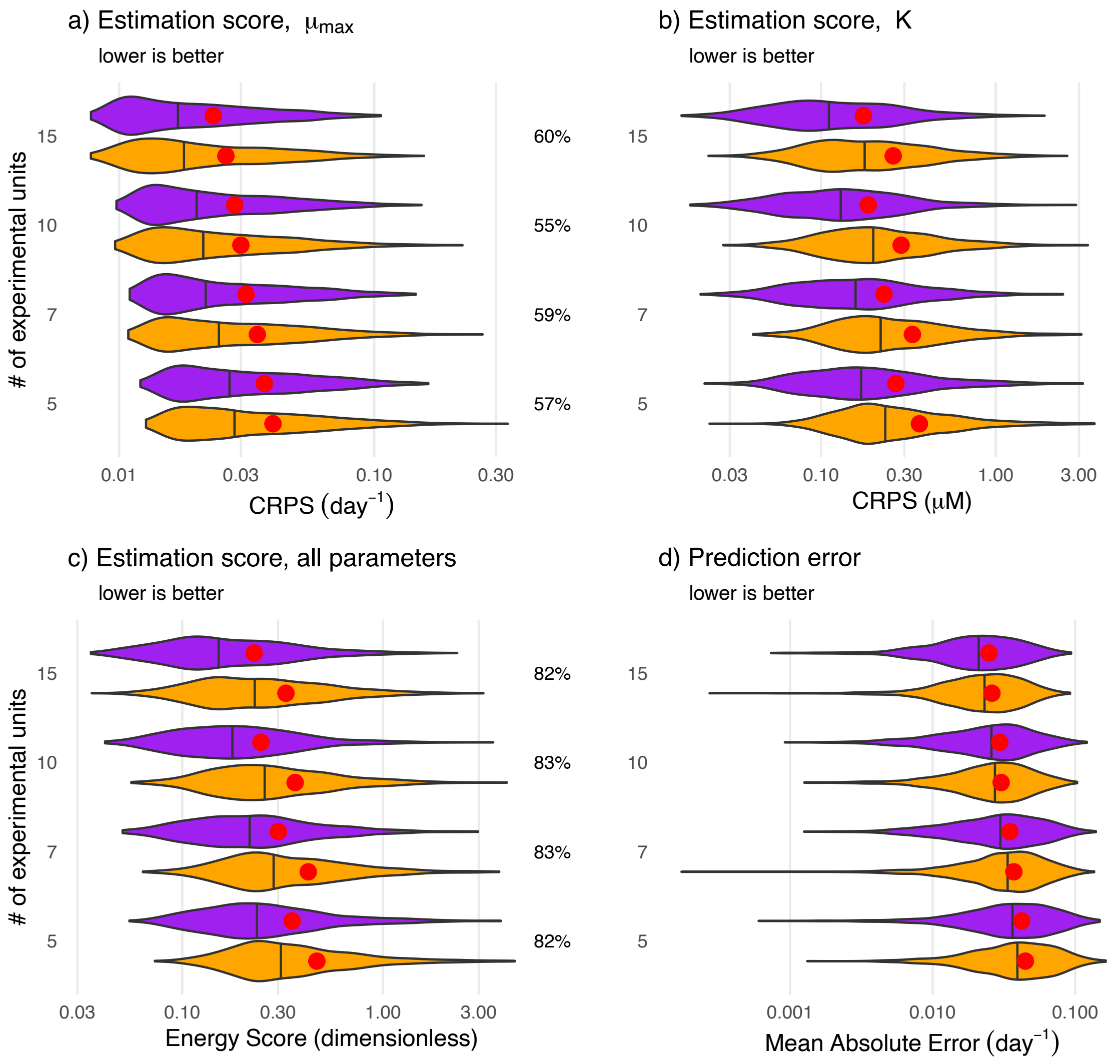

*Figure S13: Distributions of CRPS, Energy Scores and prediction errors for the Monod function, shown on a log scale. The optimal and uniform designs' scores are shown in purple and orange respectively. Vertical lines within each distribution represents its median and the red dots represent its mean. The percent of simulations where the optimal design was better is shown on the right of each plot. Optimal designs are consistently better for estimating $K$ and joint estimation of all parameters. Optimal designs are slightly better for estimating $\mu_{max}$ and aggregate prediction error.*

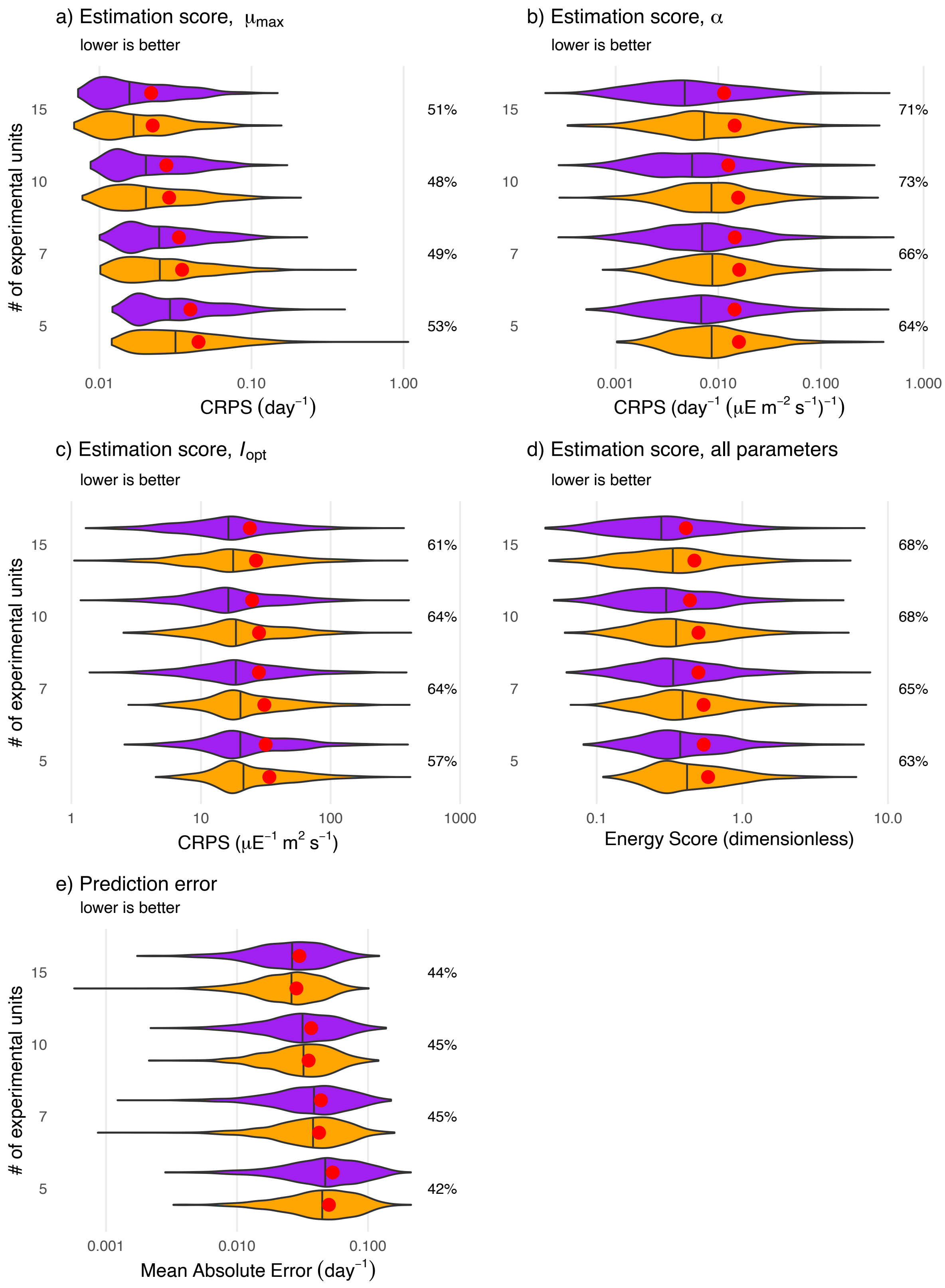

*Figure S14: Distributions of CRPS, energy scores and prediction errors for the Eilers-Peeters function shown on a log scale. The optimal and uniform designs' scores are shown in purple and orange respectively. Vertical lines within each distribution represents its median and the red dots represent its mean. The percent of simulations where the optimal design was better is shown on the right of each plot. Optimal designs are substantially better for estimating α*

*and joint estimation of all parameters. Optimal designs are slightly better for estimating* $I_{opt}$, $\mu_{max}$ *and aggregate prediction errors.*

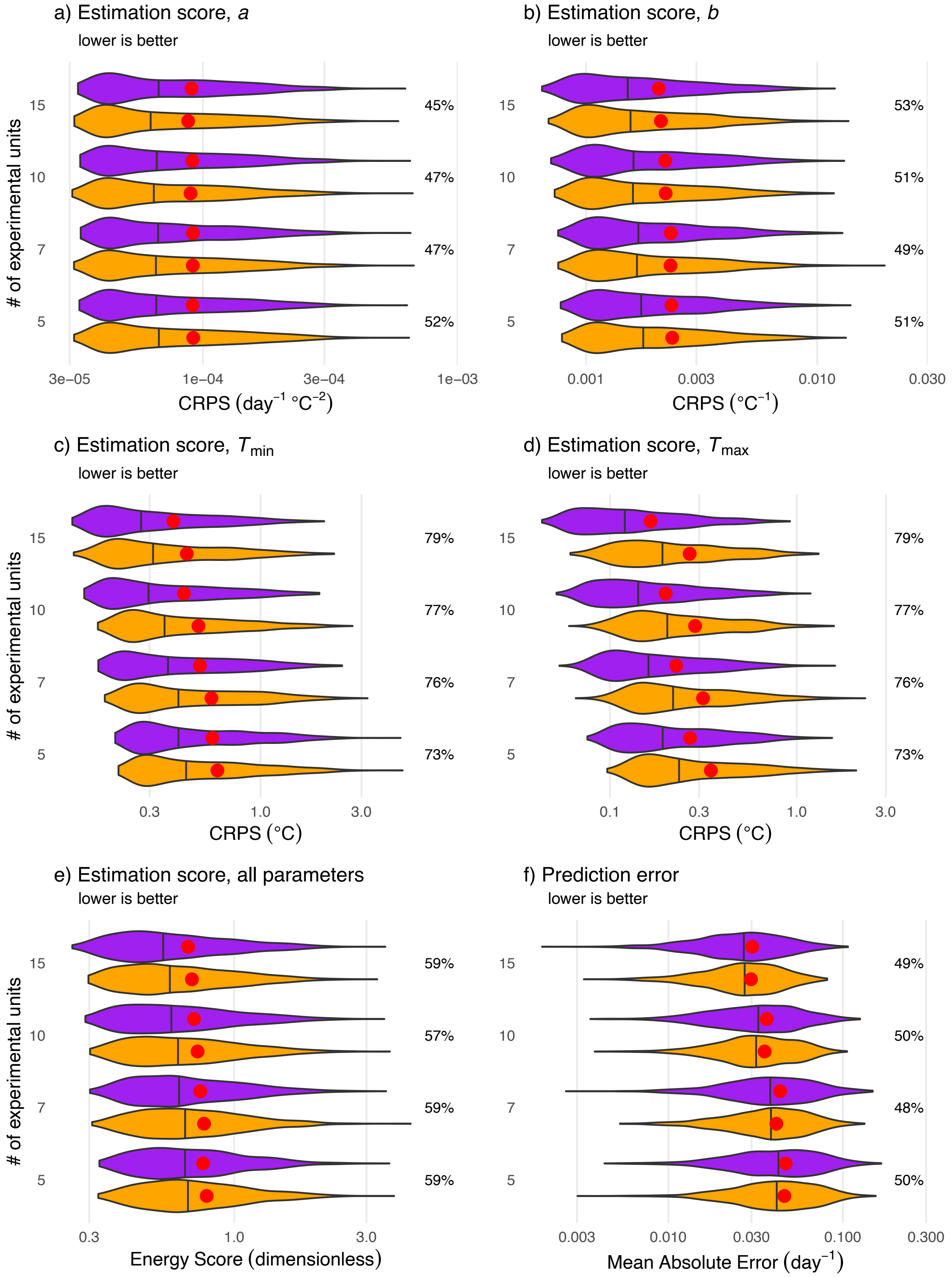

*Figure S15: Distributions of CRPS, energy scores and prediction errors for Norberg shown on a log scale. The optimal and uniform designs' scores are shown in purple and orange respectively. Vertical lines within each distribution represents its median and the red dots represent its mean. The percent of simulations where the optimal design was better is shown on the right of each plot. Optimal designs are substantially better for estimating $T_{max}$, $T_{min}$ and joint estimation of all parameters. Optimal designs are similar to uniform designs for estimating $b$ and slightly worse for estimating $a$.*

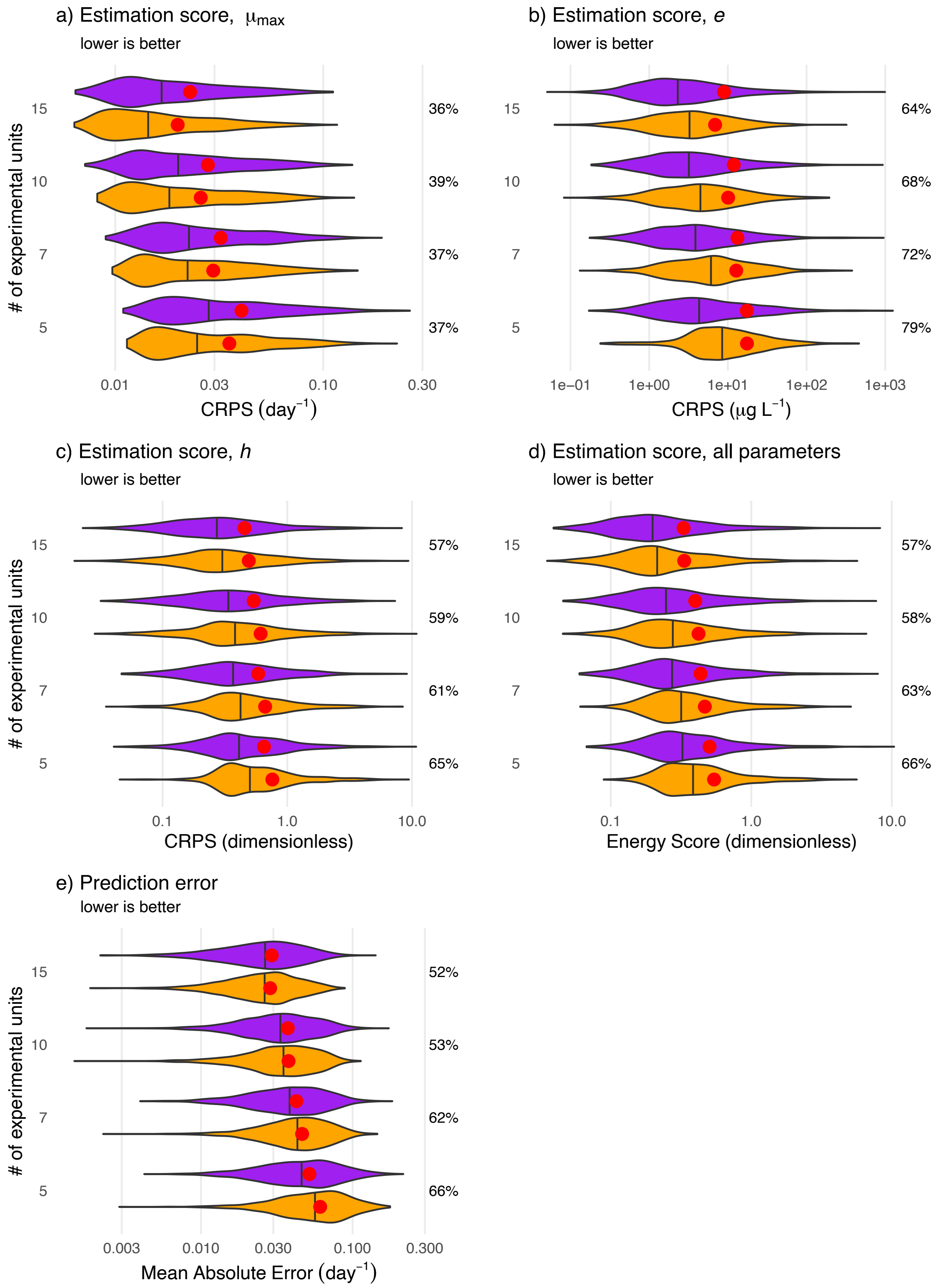

*Figure S16: Distributions of CRPS, energy scores and prediction errors for log-logistic parameters shown on a log scale. The optimal and uniform designs' scores are shown in purple and orange respectively. Vertical lines within each distribution represents its median and the red dots represent its mean. Optimal designs are substantially better*

*for estimating $e$, $h$, joint estimation of all parameters and aggregate prediction error. Optimal designs are worse for estimating $\mu_{max}$.*

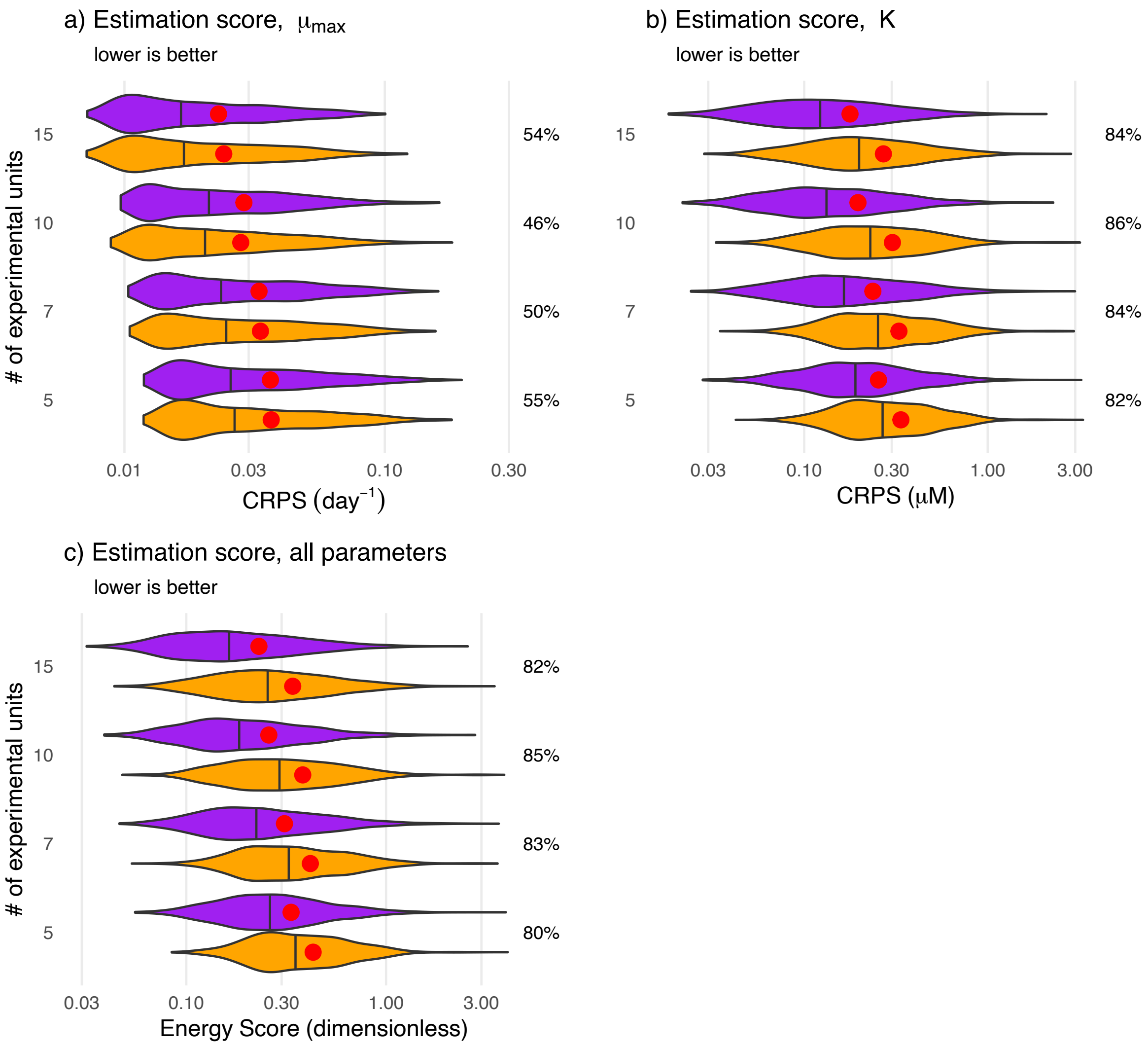

*Figure S17: Distributions of CRPS, energy scores and prediction errors for Monod parameters with bad priors shown on a log scale. The optimal and uniform designs' scores are shown in purple and orange respectively. Vertical lines within each distribution represents its median and the red dots represent its mean. Even with bad priors, optimal designs are consistently better for estimating $K$ and joint estimation of all parameters. Optimal designs are slightly better for estimating $\mu_{max}$.*

#### 8. Pointwise prediction error

Besides the aggregate prediction error shown in the main text, we also plotted prediction error at every point in a finely-spaced grid across the entire design space for all four functions. Figures S23-26 show the pointwise prediction error for Monod, Eilers-Peeters, Norberg and the log-logistic function.

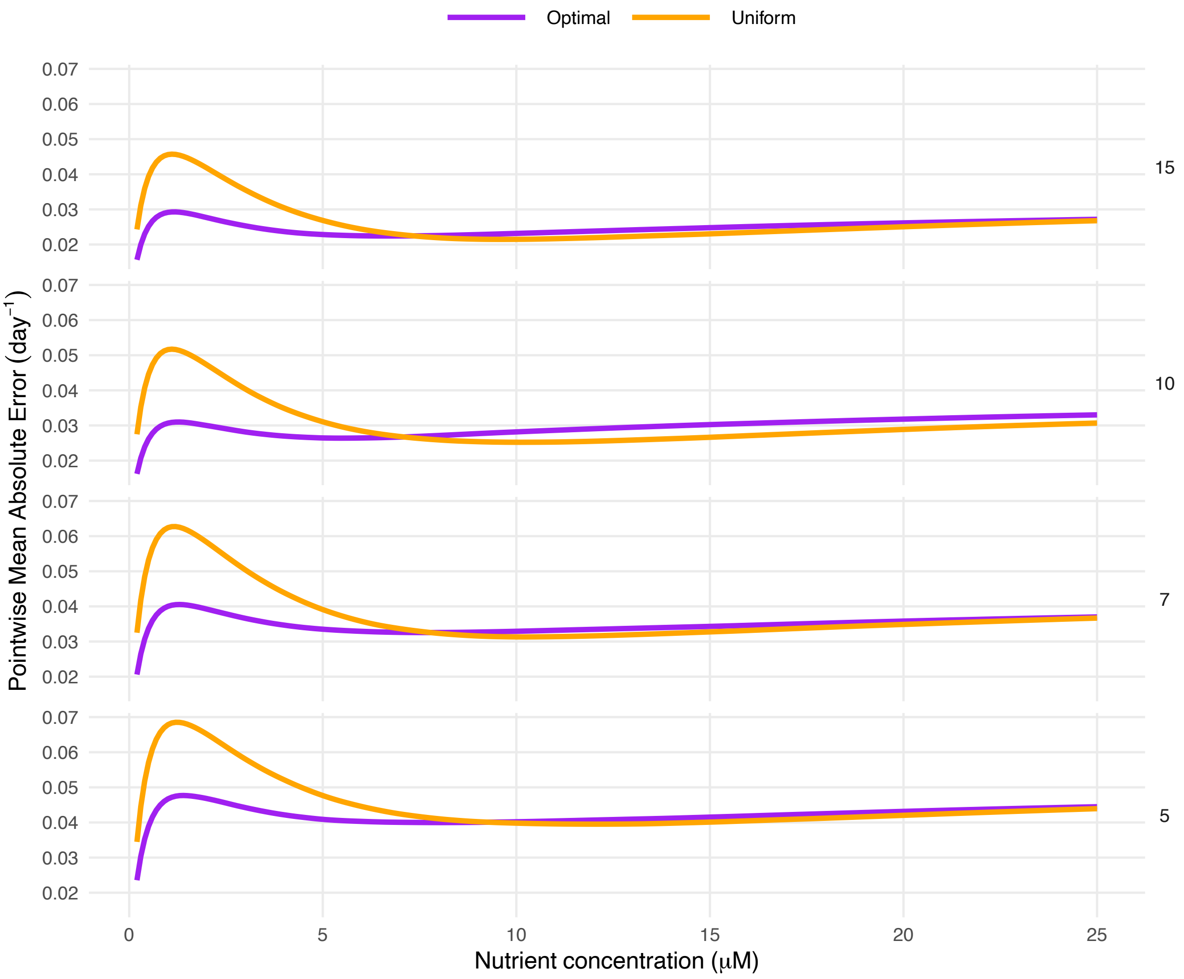

*Figure S18: Pointwise prediction error for the Monod function at 5, 7, 10 and 15 experimental units. At low nutrient concentrations, the optimal design has substantially lower prediction error than the uniform design. At high nutrient concentrations, the uniform design has slightly lower prediction errors.*

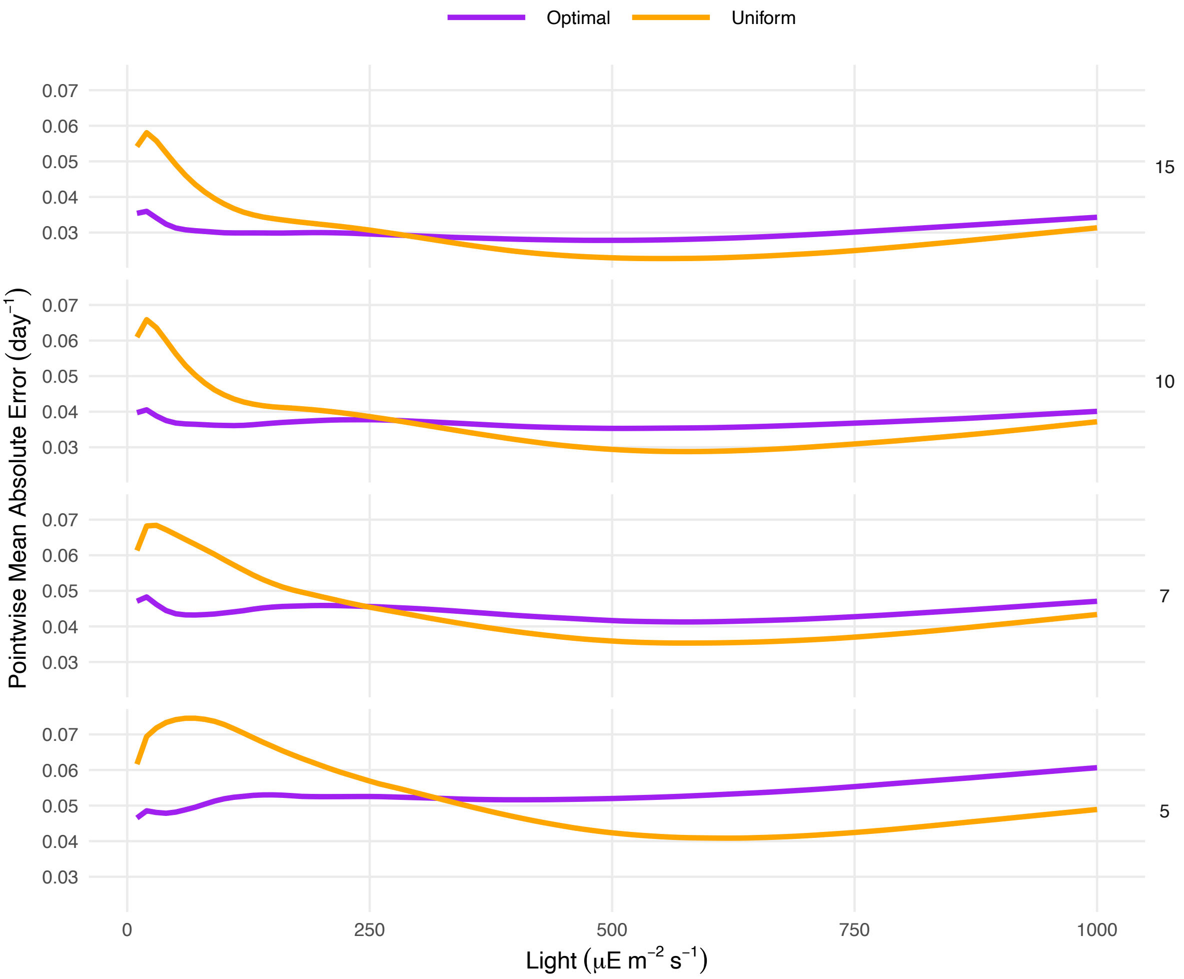

*Figure S19: Pointwise prediction error for the Eiler-Peeters function at 5, 7, 10 and 15 experimental units. At low light intensities, the optimal design has substantially lower prediction error than the uniform design. At high light intensities, the uniform design has lower prediction errors.*

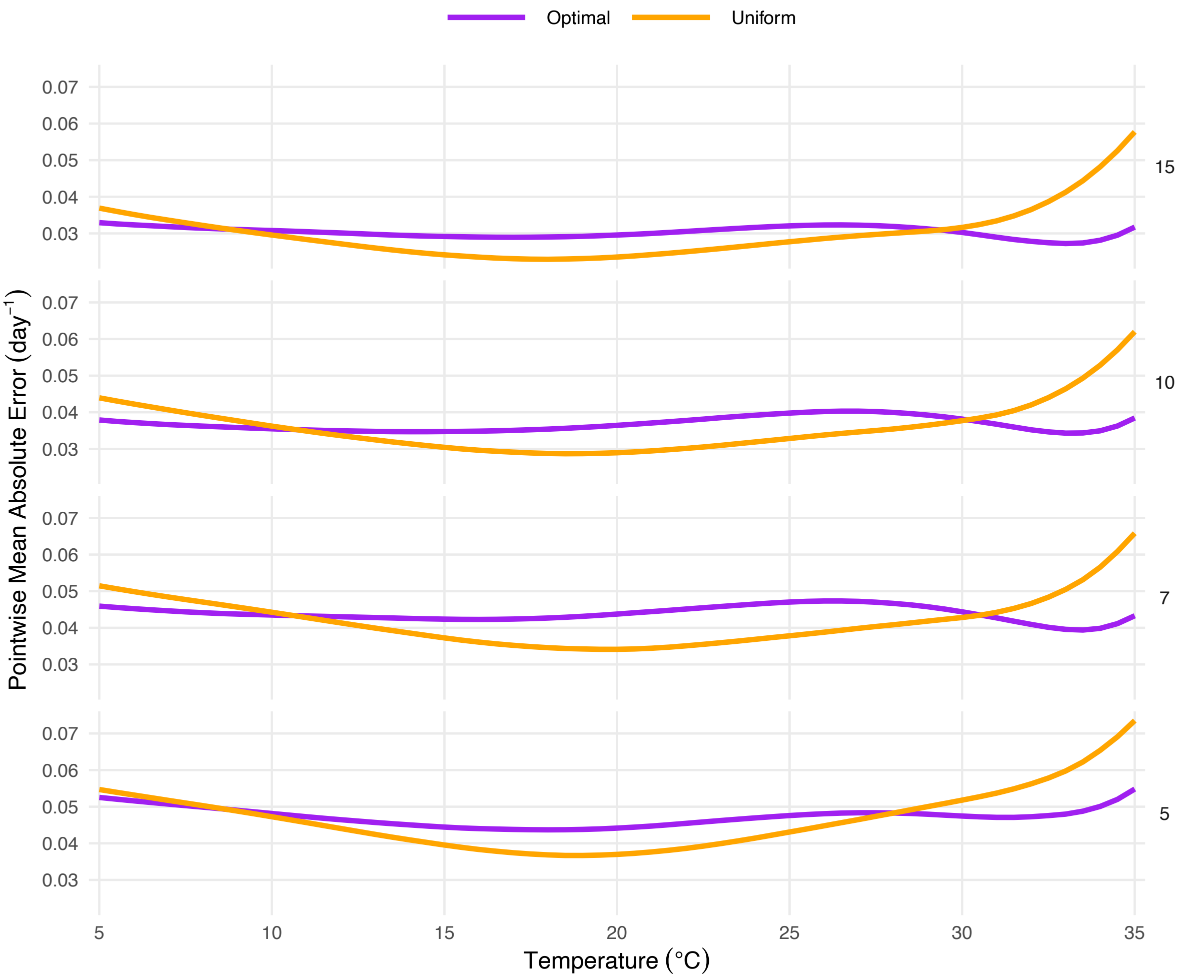

*Figure S20: Pointwise prediction error for the Norberg function at 5, 7, 10 and 15 experimental units. The optimal design has lower prediction errors at the extremes of the design space, especially near $T_{max}$, while the uniform design has lower prediction error at the center.*

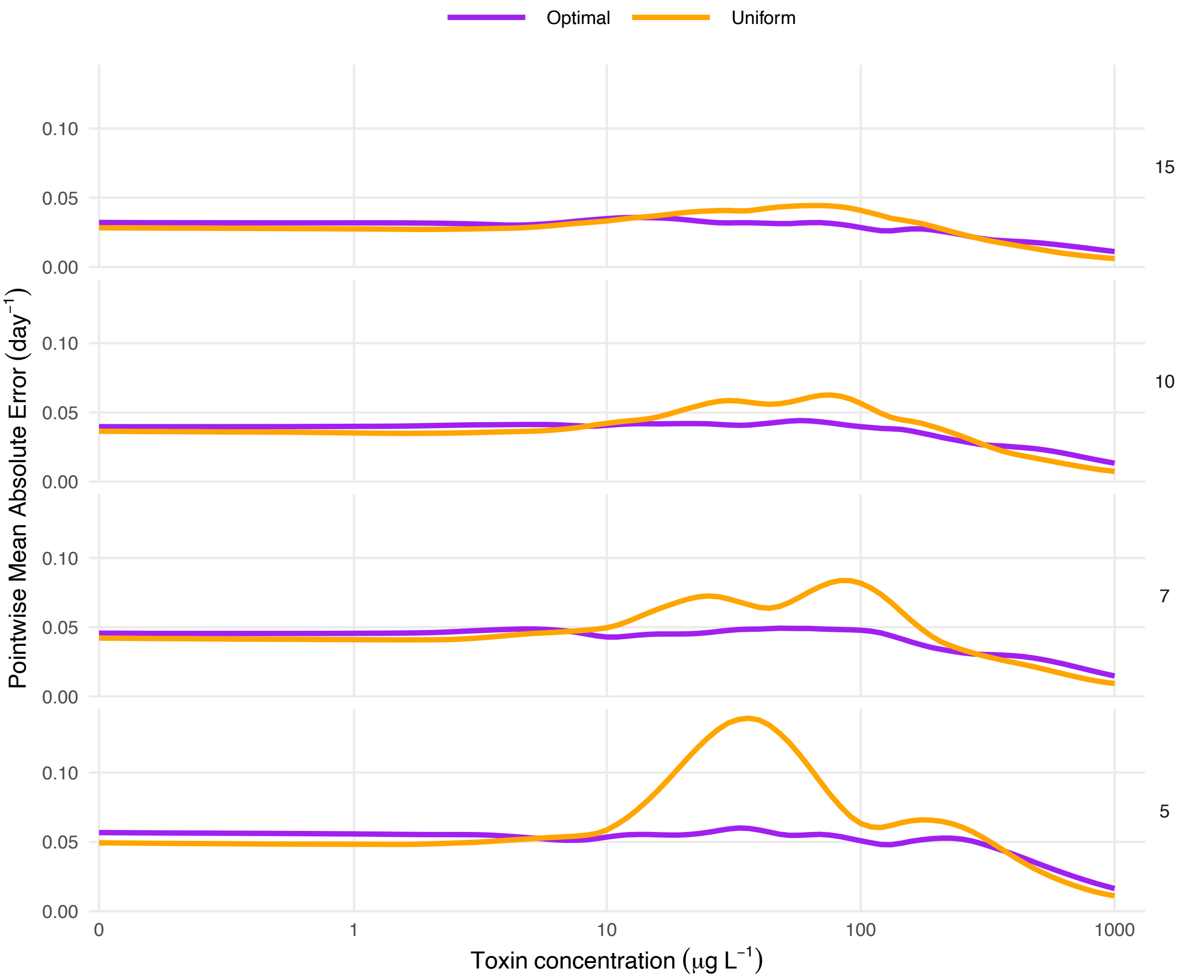

*Figure S21: Pointwise prediction error for the log-logistic function at 5, 7, 10 and 15 experimental units. At extremely low and high toxin concentrations, the uniform design has slightly lower prediction error. At low-to-intermediate toxin concentrations, the optimal design has substantially lower prediction errors.*

### 9. Finding a central curve

We generated median curves from the posteriors for each of the four functions to calculate prediction errors and from the priors to show representative curves in Figs. 2a, 3a, 4a and 5a in the main text. We used Modified Band Depth (MBD) to identify these curves. MBD is a functional data analysis technique used to identify a central curve among an ensemble of curves. Very simply, it evaluates the centrality by calculating the proportion of the design space where a curve is bounded within bands formed by other curves. It is necessary to use a technique like MBD in the context of our study since the four functions are non-linear, often with joint covariances among the parameters. Thus, simpler methods like just using the means/medians of the prior distributions does not translate into a mean/median curve. MBD guarantees that the selected median curve is from an actual parameter draw from the prior/posterior and is therefore both central and biologically realistic. The parameters corresponding to the median curve in Figs. 2a, 3a, 4a and 5a in the main text are provided below. For more details about MBD, see López-Pintado & Romo, 2009.

*Table S1: Parameter values corresponding to the median curves identified through MBD from the prior distributions and shown in Figs. 2a, 3a, 4a and 5a in the main text. All values are rounded to four decimal places.*

| Function | Parameter | Value corresponding to median curve from the priors | Units |
|---|---|---|---|
| Monod | $\mu_{\max}$ | 1.0493 | $\mathrm{day}^{-1}$ |
| | $K$ | 1.5185 | $\mu\mathrm{M}$ |
| Eilers-Peeters | $\mu_{\max}$ | 0.9690 | $\mathrm{day}^{-1}$ |
| | $\alpha$ | 0.0496 | $\mathrm{d}^{-1}\ (\mu\mathrm{E\ m}^{-2}\ \mathrm{s}^{-1})^{-1}$ |
| | $I_{\mathrm{opt}}$ | 226.1083 | $\mu\mathrm{E\ m}^{-2}\ \mathrm{s}^{-1}$ |
| Norberg | $a$ | 0.0018 | $\mathrm{d}^{-1}\ {}^{\circ}\mathrm{C}^{-2}$ |
| | $b$ | 0.0284 | ${}^{\circ}\mathrm{C}^{-1}$ |
| | $T_{\max}$ | 34.8837 | ${}^{\circ}\mathrm{C}$ |
| | $T_{\min}$ | 1.5899 | ${}^{\circ}\mathrm{C}$ |
| Log-logistic | $\mu_{\max}$ | 1.0084 | $\mathrm{day}^{-1}$ |
| | $e$ | 47.3726 | $\mu\mathrm{G\ L}^{-1}$ |
| | $h$ | 2.2599 | |
